# Multi-factorial regulatory networks of placental antibody transfer by Fc receptors and maternal IgG Fc characteristics are modulated by clinical covariate profiles

**DOI:** 10.64898/2026.05.06.723089

**Authors:** Remziye E. Wessel, Anna Sawik, Abigail Boyette, Guadalupe Martinez, Vanessa Kirschner, Caitlin Sullivan, Xu Yang, Lisa Zuckerwise, Parastoo Azadi, Ileana S. Mauldin, Donald J. Dudley, Yalda Afshar, Sepideh Dolatshahi

**Affiliations:** Department of Biomedical Engineering, University of Virginia School of Medicine, Charlottesville, Virginia 22908; Beirne B. Carter Center for Immunology Research, University of Virginia School of Medicine, Charlottesville, Virginia 22908; Division of Maternal-Fetal Medicine, Department of Obstetrics and Gynecology, University of California, Los Angeles, Los Angeles, California 90095; Complex Carbohydrate Research Center, University of Georgia, 315 Riverbend Road, Athens, Georgia 30602; Division of Maternal-Fetal Medicine, Department of Obstetrics and Gynecology, University of Virginia School of Medicine, Charlottesville, Virginia 22908; Department of Surgery, University of Virginia School of Medicine, Charlottesville, Virginia 22908; Molecular Biology Institute, University of California, Los Angeles, Los Angeles, California 90095

## Abstract

Maternal immunoglobulin G (IgG) transferred across the placenta is crucial for newborn immunity. IgG transfer efficiency modulated by Fc characteristics including subclass and glycosylation gives rise to diverse transfer profiles across the population, yet the molecular mechanisms driving this variation are incompletely understood. To disentangle multimodal molecular relationships driving population heterogeneity in maternal-fetal antibody transfer, we characterized placental and serological antibody features in matched human tissues from two geographically distinct cohorts. Unsupervised clustering of maternal clinical covariates in both cohorts revealed a distinct patient profile with reduced plasma C-reactive protein and pregravid BMI and enhanced transfer efficiency of IgG subclasses. Quantification of IgG Fc glycan structures in matched maternal and cord plasma by liquid chromatography-mass spectrometry revealed subclass-specific IgG glycosylation patterns which impacted placental transfer efficiency and correlated with these key clinical features. We profiled expression and colocalization of key Fc γ Receptors (FcγRs) by multiplex immunohistochemistry, revealing cell type-specific expression patterns. Variable FcγR expression across gestation was consistent in both cohorts, implicating FcγRs as key drivers of temporal antibody transfer dynamics. While FcγRs were not strongly variable across the clinical profiles, partial correlation analysis of matched samples controlling for gestational age and demographic covariates revealed correlations between FcγR expression frequencies and glycan- and subclass-specific transfer efficiency. These data systematically define multi-factorial regulatory networks of antibody transfer by placental Fc receptors and maternal IgG Fc characteristics, which are further modulated by clinical covariates. This study provides a basis for the rational design of prenatal vaccination strategies, administration schedules, and potential lifestyle interventions to improve maternal-fetal immunity.

## Introduction

Maternal immunoglobulin G (IgG) transferred across the placenta is critical for providing newborns with humoral immune protection against pathogenic threats. Beyond neutralization, maternal antibodies prime both innate and adaptive responses during primary pathogen exposures in early-life^1^ and exert long-term impacts on the offspring’s B cell repertoire and recall response strength into adulthood^2^, underscoring the importance of maternal antibodies in human immune development. The placenta transfers IgG in an Fc-selective manner, generally resulting in efficient transfer of IgG1, IgG3, and IgG4 (but not IgG2) and digalactosylated IgG^3–7^. However, several studies have reported nuanced differences in IgG transfer efficiency across cohorts with respect to subclass-, glycan-, and antigen-specificity^3^. The underlying causes of these cross-cohort differences are unknown but may point toward multifaceted regulation of antibody transfer by placental tissue and maternal IgG Fc characteristics^8^.

While the role of FcRn in IgG transcytosis in syncytiotrophoblasts (STBs) is undisputed, other FcγRs expressed by placental STBs and microvascular endothelial cells (ECs)—especially FcγRIIIa and FcγRIIb—have recently gained attention as possible additional regulators of antibody transfer^9^. FcRn is a high-affinity receptor for IgG that does not bind preferentially to different Fc glycoforms, whereas FcγRIIIa and FcγRIIb binding affinity varies with subclass and Fc glycosylation; this variation may drive Fc-selective antibody transfer^5^. While a few prior studies have bolstered this hypothesis by uncovering associations between FcγR expression in human placental tissue with subclass- and glycan-specific IgG transfer efficiency^10,11^ and identifying *in situ* colocalization between FcRn and FcγRs^4,5,10^, these multimodal relationships have not been systematically characterized. Moreover, mechanistic evidence that FcRn is sufficient for transcytosis in STBs and ECs calls into question the extent to which these FcγRs regulate transfer^12,13^, and whether they are involved with the canonical pathway driven by FcRn.

Cord IgG titers rise steadily throughout pregnancy; the rate of transfer increases exponentially approaching parturition, when cord IgG levels typically exceed maternal concentrations^14^. Whether the apparent increase in transfer efficiency is due to increased placental volume, changes in Fc receptor expression^8^, or simply reflects a gradual accumulation of IgG throughout pregnancy is unclear. Previous work has revealed similar functional profiles of antibodies in preterm and term neonates despite lower absolute IgG titers against several antigen-specific subtypes^15^. Others have shown that preterm newborns have decreased frequency of IgG Fc sialylation and an increased proportion of agalactosylated IgG, both of which are associated with pro-inflammatory functionality^16,17^. Together, these findings suggest Fc-dependent placental antibody transfer is temporally regulated, resulting in early transfer of potent pro-inflammatory antibodies capable of engaging diverse immune effector functions. Therefore, there may exist a gestationally-defined window of opportunity to administer prenatal vaccines which yields both maximum absolute antibody titers and functional benefit for newborns.

To decipher networks regulating placental antibody transfer, we systematically probed plasma antibody Fc characteristics and placental Fc receptor expression in two geographically distinct U.S.-based cohorts. Unsupervised analysis of patient metadata revealed distinct maternal profiles associated with variable subclass transfer efficiency. Our data indicate the variation across these clusters may be driven by subclass-specific Fc glycosylation, which is further modulated by placental Fc receptor expression. Fc receptors displayed highly cell type-specific expression and tightly coordinated temporal expression trajectories, implicating FcRn and FcγRs as gatekeepers of Fc-selective transfer and rate-determining features in transfer dynamics across gestation. Ultimately, our study identifies multimodal regulatory networks which may contribute to heterogeneity in antibody transfer efficiency across the population, providing a basis for precision prenatal vaccines to tune and personalize antibody transfer for optimal neonatal immunity.

## Results

### Maternal clinical and demographic features define distinct IgG subclass transfer profiles in two independent sample populations

IgG transfer efficiency is highly variable across published studies and varies with Fc characteristics including subclass and Fc glycosylation^3^. We hypothesized that this heterogeneity is driven by a combination of maternal immune features dictating IgG Fc quality and placental Fc receptors, which selectively bind to different Fc glycoforms and subclasses with variable affinity^18^. To disentangle the relative contribution of maternal and placental features governing antibody transfer efficiency, we systematically analyzed maternal and umbilical cord IgG Fc features with matched placental tissue from patients in two geographically distinct cohorts spanning a range of gestational ages (**Figure 1A-B**). Cohort 1 comprised a total of 100 individuals spanning first, second, and third trimester (see Methods) (**Table 1**). Cohort 2, which served as a validation cohort, comprised only live births from third trimester containing a mixture of preterm and term specimens (n=45) (**Table 2**). Participants had either formalin-fixed, paraffin-embedded (FFPE) placental tissue available, maternal and umbilical cord plasma, and a subset from Cohort 1 (n=27) and Cohort 2 (n=10) had matched placental and plasma specimens (**Figure 1C**). Patients in both cohorts displayed a broad range of IgG subclass transfer efficiencies, but subclass-specific transfer was similarly variable in both cohorts spanning nearly two orders of magnitude (**Figure 1D**).

**Figure 1.**
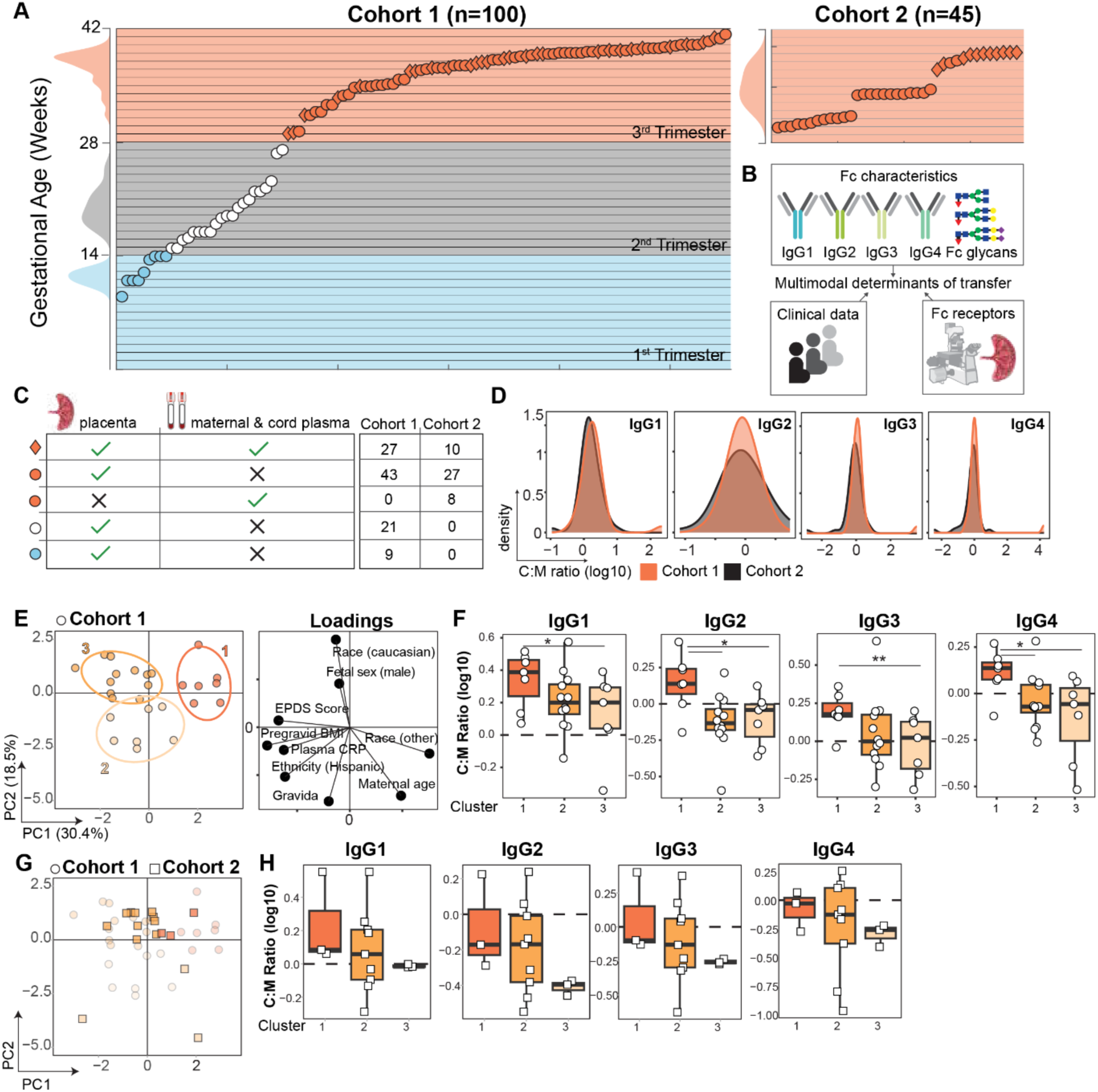
Unsupervised analysis reveals clinical signatures IgG subclass transfer variability. *(A)* Scatter plot showing the distribution of gestational ages in Cohort 1 (left) and Cohort 2 (right). The density plots on the left axis of each plot show the distribution of gestational ages in each trimester group (first trimester, blue; second trimester, white/gray; third trimester, orange). *(B)* Schematic of the data collected in this study. A combination of Fc characteristics, clinical data, and placental Fc receptors and relationships between features were analyzed to identify multimodal determinants of transfer. Clinical data, IgG isotypes, and Fc receptors were measured in both cohorts. Fc glycans were measured in a subset of patients from Cohort 2. (C) Table summarizing the available sample types. Patients for whom both FFPE placental tissue and paired maternal and umbilical cord plasma were available are represented by diamonds (n=37). Patients for whom only FFPE placental tissue (n=100) or only maternal and cord plasma (n=8) was available are represented by circles. Points are colored according to trimester of delivery or termination. The number of patients in each category is tabulated for each cohort on *the right.* (D) Density plots show the distribution of IgG subclass C:M ratios in each cohort (n=27, Cohort 1; n=18, Cohort 2). Data are log10-transformed such that C:M ratio of zero corresponds to equal concentrations in maternal and cord plasma. (E) Right, PCA scores projection reducing nine clinical variables into 2D space from Cohort 1 patients with plasma samples and complete metadata available (n=26). The percent variance captured by PC1 and PC2 is provided on the axes (49% total variance). Each point represents one patient profile. Points are colored by k-means cluster identity, corresponding to the ellipses showing 75% confidence intervals for each cluster. Left, biplot shows feature loadings on PC1 and PC2. (F) Boxplots indicate the average IgG subclass cord:maternal ratios across clusters. The dashed line at log10(C:M ratio)=0 indicates equal concentrations in cord and maternal plasma, and log10(C:M ratio)>0 denotes efficient antibody transfer. P-values shown are from a Kruskal-Wallis test with Dunn’s test for multiple comparisons (*p<0.05, **p<0.01). (G) Patient profiles from Cohort 2 with plasma samples and complete metadata available (n=16) were projected into the PC space defined by Cohort 1 by multiplying each patient profile by feature loadings in (E). Each patient was assigned a cluster corresponding to those in (E) based on the minimum Euclidean distance to the cluster centroids. Faint circles from Cohort 1 are overlaid for reference. (H) Boxplots indicate the average IgG subclass cord:maternal ratios across clusters. The dashed line at log10(C:M ratio)=0 indicates equal concentrations in cord and maternal plasma, and log10(C:M ratio)>0 denotes efficient antibody transfer. No comparisons were statistically significant at p<0.05 (Kruskal-Wallis test with Dunn’s test for multiple comparisons). PCA, principal components analysis. EPDS, Edinburgh Postnatal Depression Scale. CRP, C-reactive protein.

**Table 1.**
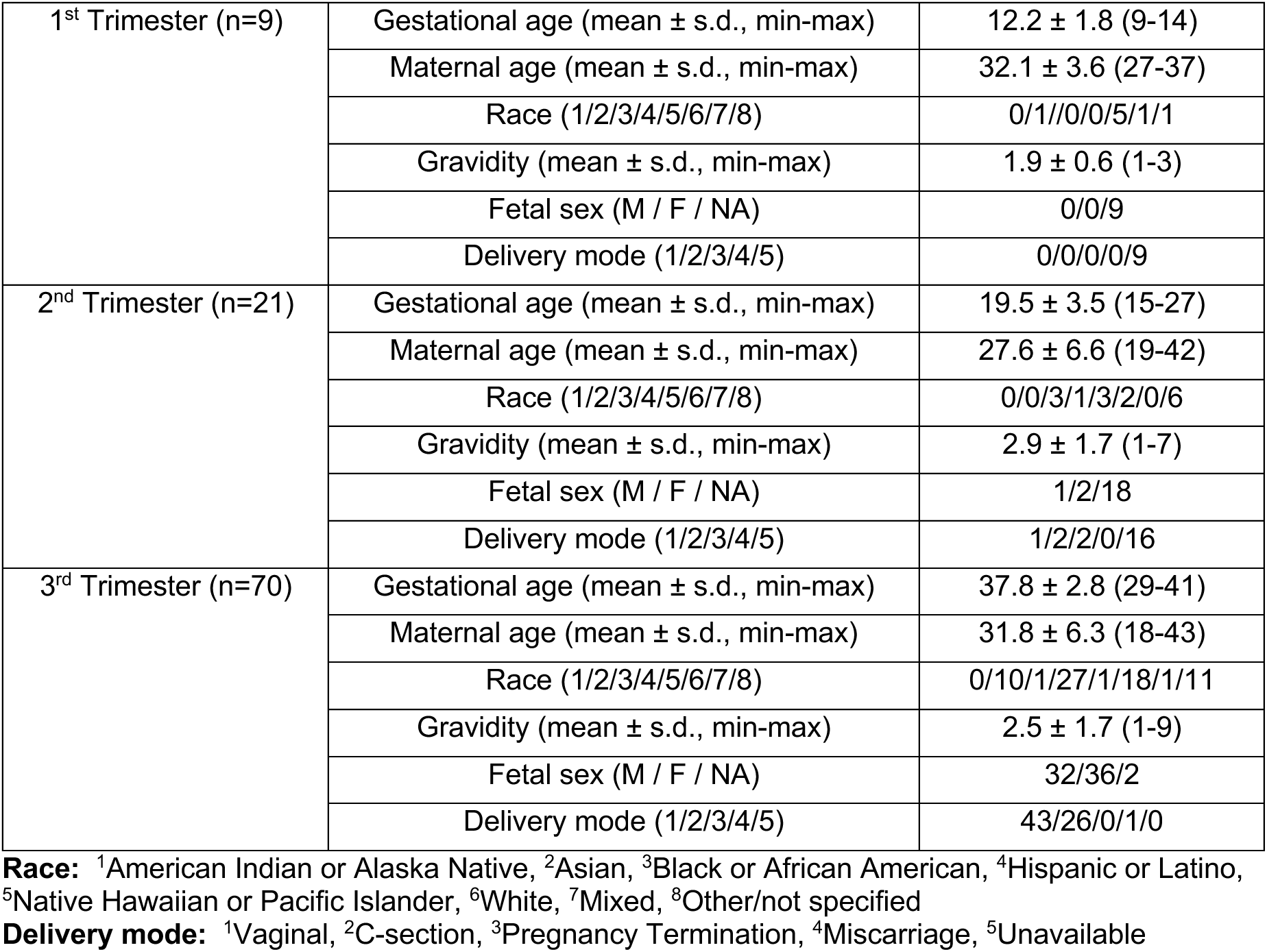
Cohort 1 characteristics.

**Table 2.**
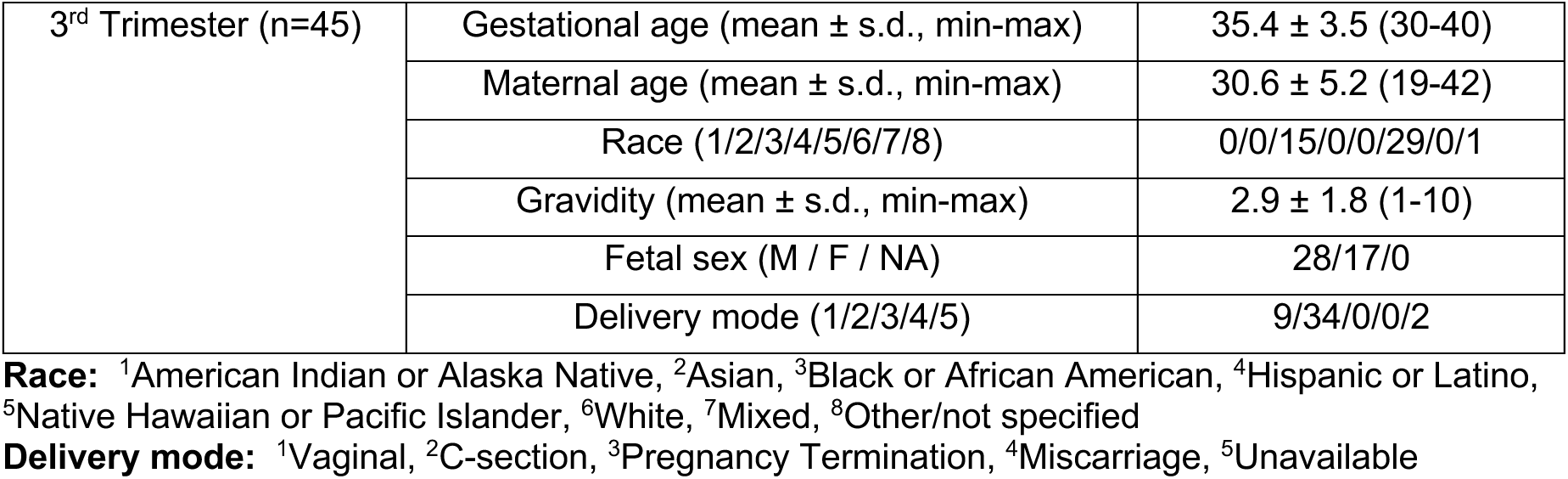
Cohort 2 characteristics.

|  |  |  |
| --- | --- | --- |
| 3<sup>rd</sup> Trimester (n=45) | Gestational age (mean ± s.d., min-max) | 35.4 ± 3.5 (30-40) |
|  | Maternal age (mean ± s.d., min-max) | 30.6 ± 5.2 (19-42) |
|  | Race (1/2/3/4/5/6/7/8) | 0/0/15/0/0/29/0/1 |
|  | Gravidity (mean ± s.d., min-max) | 2.9 ± 1.8 (1-10) |
|  | Fetal sex (M / F / NA) | 28/17/0 |
|  | Delivery mode (1/2/3/4/5) | 9/34/0/0/2 |

**Race:** <sup>1</sup>American Indian or Alaska Native, <sup>2</sup>Asian, <sup>3</sup>Black or African American, <sup>4</sup>Hispanic or Latino, <sup>5</sup>Native Hawaiian or Pacific Islander, <sup>6</sup>White, <sup>7</sup>Mixed, <sup>8</sup>Other/not specified
**Delivery mode:** <sup>1</sup>Vaginal, <sup>2</sup>C-section, <sup>3</sup>Pregnancy Termination, <sup>4</sup>Miscarriage, <sup>5</sup>Unavailable

To define profiles of patient-intrinsic features which may underscore this cross-cohort variability, we performed unsupervised clustering of clinical metadata. Principal components analysis (PCA) on clinical data from Cohort 1 captured 49% of the variance on the first two PCs (**Figure 1E**), and patients separated into three distinct clusters in this latent space. Cluster 1 patients displayed a purportedly anti-inflammatory phenotype, characterized by low pregravid BMI, low plasma CRP (a biomarker of systemic inflammation), and low Edinburgh Postnatal Depression Scale (EPDS) scores, a quantitative metric based on a 10 question survey used as a proxy for psychosocial stress (high EPDS score indicates greater stress)^19,20^ (**Figure 1E, Supplemental Figure S1**). These patients were entirely of non-Hispanic, non-Caucasian ethnicity and tended to have higher maternal age at birth. Conversely, patients in Clusters 2 and 3 tended to have higher plasma CRP, higher EPDS scores, and higher pregravid BMI, together indicating an inflammatory state. All four IgG subclasses displayed variable transfer efficiency across the clusters, with all subclasses displaying enhanced transfer efficiency in Cluster 1 compared to Cluster 3 (*p*<0.05) (**Figure 1F**). To determine whether these patient-specific transfer profiles were generalizable in Cohort 2, we projected the subset of patient profiles from Cohort 2 with complete metadata and IgG subclass measurements into this latent space defined by Cohort 1 (Methods). Each patient profile from Cohort 2 was assigned a cluster identity based on the minimum Euclidean distance to the cluster centroids in Cohort 1 (**Figure 1G**). This revealed a similar trend of lower transfer efficiency across all IgG subclasses in Cluster 3 (not significant) (**Figure 1H**). Therefore, maternal clinical and demographic profiles partially explain variation in IgG subclass transfer, and metabolic and social stress and inflammation may negatively impact placental antibody transfer.

### Subclass-specific Fc glycans underscore inter-patient antibody transfer heterogeneity

IgG carries a conserved N-linked glycosylation site at Asn297 on each heavy chain of the Fc region (**Figure 2A**). The majority of serum IgG in humans is decorated with complex glycans which exert strong effects on IgG binding to Fc receptors and downstream effector functions^21,22^. Therefore, we posited that maternal clinical covariates affect IgG subclass-specific transfer efficiency by modulating maternal IgG Fc glycosylation, which in turn affects FcγR binding and placental transfer^4–6^. To determine the effect of Fc glycans on subclass-specific IgG transfer, we performed subclass-specific IgG glycoproteomics in a subset of mother and cord dyads (n=14 pairs) from Cohort 2 (**Supplemental Figure S2, Supplemental Tables S1-2**). We found that terminal sialylation and digalactosylation enhanced transfer of all IgG subclasses, with cord levels displaying an enrichment of these glycoforms relative to maternal levels (**Figure 2B-C**). Fucosylated, digalactosylated IgG lacking terminal sialic acid (G2F) displayed efficient transfer for IgG2/3, but not for IgG1 and IgG3/4 (**Supplemental Figure S2**). Meanwhile, bisecting N-Acetylglucosamine (GlcNAc) structures exerted subclass-specific effects on transfer efficiency, with bisected IgG2/3 displaying inefficient transfer and bisected IgG3/4 significantly higher in cord relative to maternal plasma (**Figure 2B-C**). Monogalactosylated IgG1 and IgG2/3 were reduced in cord plasma compared to maternal plasma, but there was no difference detected in the IgG3/4 cluster. Fc sialylation was thus universally associated with increased placental transfer, whereas bisecting GlcNAc and monogalactosylation exhibited subclass-specific effects.

**Figure 2.**
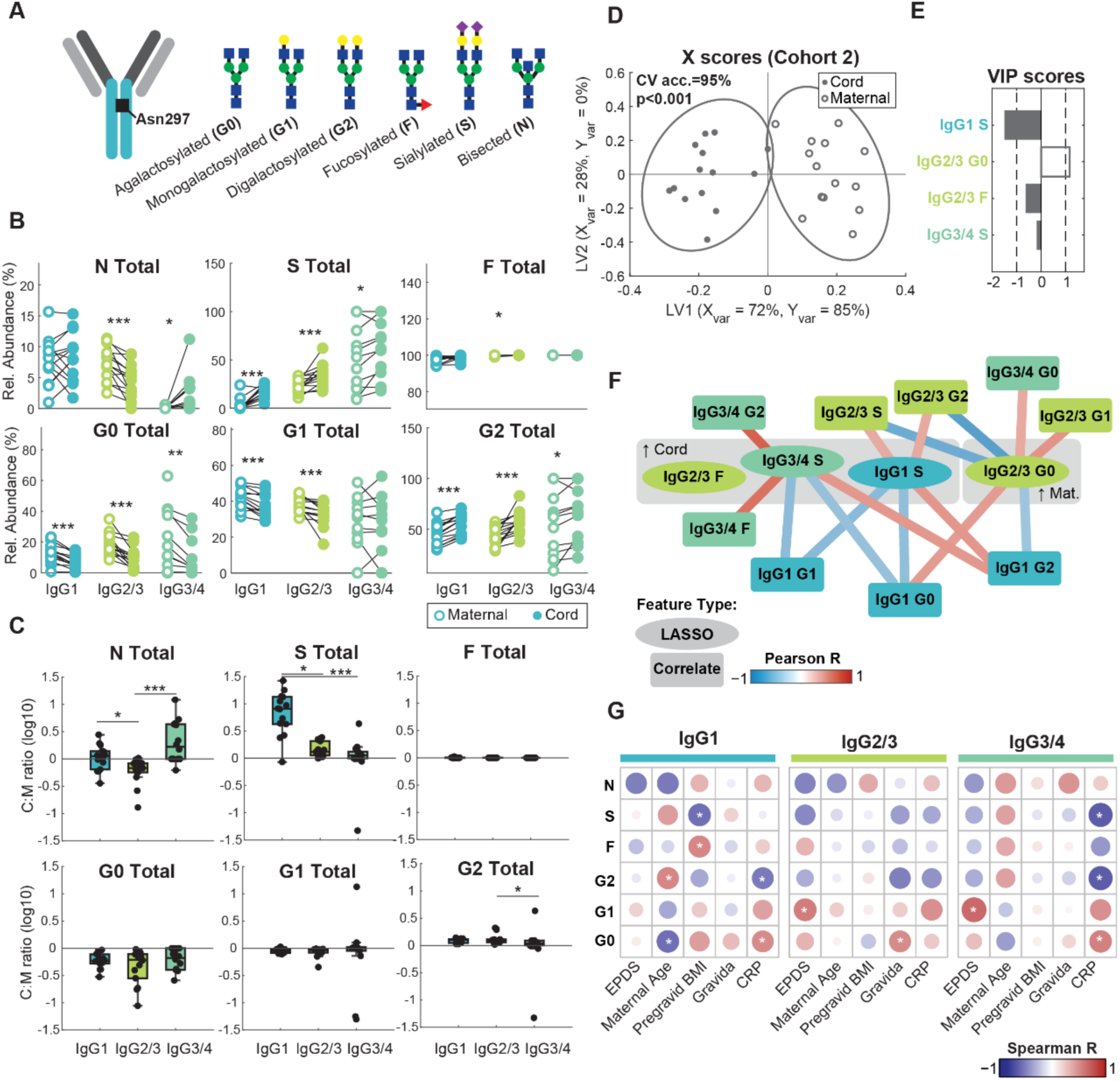
Fc glycosylation determines subclass-specific IgG transfer and is perturbed by maternal covariates. (A) Schematic of complex Fc N-glycans quantified in this study. (B) Paired analysis comparing relative abundance of summary glycan structures for three distinct IgG subclass groups (IgG1, IgG2/3, and IgG3/4) in maternal plasma (open circles) versus umbilical cord plasma (filled circles) for a subset of patients in Cohort 2 (n=14 pairs). (*p<0.05, **p<0.01, ***p<0.001, Wilcoxon signed-rank test). (C) Boxplots indicate the average glycan- and subclass-specific IgG cord:maternal ratios. The line at log10(C:M ratio)=0 indicates equal concentrations in cord and maternal plasma, and log10(C:M ratio)>0 denotes efficient antibody transfer. P-values shown are from a Kruskal-Wallis test with Dunn’s test for multiple comparisons (*p<0.05, ***p<0.001). (D) X Scores plot for PLSDA model distinguishing between maternal and umbilical cord samples based on relative abundance of subclass-specific glycosylation motifs. The variance in X and Y captured by each latent variable (LV1 and LV2) is denoted on the axis labels. Significance was determined by a two-sided t-test comparing the true model accuracy scores over 1,000 independent rounds of 5-fold cross-validation against 1,000 models trained on data with randomly permutated class labels. (E) VIP scores bar plot displays the relative importance of each LASSO-selected feature to model performance. VIP > 1 denotes features with greater than average influence on the model. VIP scores are artificially oriented and according to their loadings on LV1. Open bars correspond to features enriched in maternal plasma and filled bars correspond to features enriched in cord plasma. (F) LASSO co-correlate network displays pairwise Pearson correlations among LASSO-selected features (ovals) with other features not included in the model (rectangles). Edges are colored according to the strength of the pairwise correlations. Nodes are colored by IgG subclass. The network was trimmed to only show significant correlations (p<0.05). (G) Heatmap of Spearman correlations between continuous-valued maternal clinical covariates and relative abundance of IgG subclass-specific summary glycans in maternal plasma (n=14). Color corresponds to the correlation coefficient value, and size of the circle corresponds to significance level. Significant correlations are highlighted with an asterisk (*p<0.05). PLSDA, partial least squares discriminant analysis. CV acc., cross validation accuracy. VIP, variable importance in projection. LASSO, least absolute shrinkage and selection operator.

To identify the key subclass-specific glycoforms that differed between maternal and umbilical cord antibodies, we performed partial least squares discriminant analysis (PLSDA) with least absolute shrinkage and selection operator (LASSO) feature selection discriminating between matched maternal and cord plasma samples. The resulting PLSDA model trained on four LASSO-selected features had a mean cross-validation accuracy of 95% and was significant by permutation test (*p*<0.001) (**Figure 2D**). The features which contributed most to the separation between maternal and cord samples based on variable importance in projection (VIP) scores (VIP>1) were sialylated IgG1 and agalactosylated IgG2/3 enriched in maternal and cord antibodies, respectively (**Figure 2E**). As LASSO minimizes colinear relationships among features, we next explored relationships with other features not selected by LASSO using Pearson correlation network analysis (**Figure 2F**). IgG2/3 agalactosylation was positively correlated with agalactosylation of IgG1 and IgG3/4, suggesting consistent regulation of galactosylation across subclasses. IgG3/4 sialylation correlated with IgG3/4 fucosylation, reflecting prerequisite fucosylation for further glycan modifications. Together, these data revealed subclass-specific effects of IgG glycosylation on antibody transfer efficiency and possible subclass-specific regulation of IgG glycosylation.

IgG glycans have been shown to vary with features such as maternal age, BMI, markers of metabolic health and inflammation^23–25^. We hypothesized that the variation in IgG transfer efficiency across clusters observed in **Figure 1** was driven in part by perturbations to IgG glycosylation with these clinical covariates. To test this hypothesis, we performed pairwise Spearman correlation analysis between key clinical covariates and summary Fc glycans. For instance, maternal age was positively correlated with IgG1 digalactosylation frequency, consistent with increased IgG1 transfer efficiency observed in the patient cluster with high maternal age (**Figure 2F, Supplemental Figure S1**). On the other hand, maternal CRP was inversely associated with IgG1 and IgG3/4 digalactosylation and IgG3/4 sialylation, and pregravid BMI was inversely associated with IgG1 sialylation (*p*<0.05). Both CRP and BMI were elevated in clusters with diminished IgG transfer efficiency (**Figure 1E-H, Supplemental Figure S1**), suggesting that these inflammatory perturbations alter the glycosylation profile in a manner not conducive to efficient transfer. EPDS score was positively associated with IgG2/3 and IgG3/4 monogalactosylation (*p*<0.05) but negatively associated with digalactosylation (not significant). These relationships were partially validated in a second batch of samples from pregnant mothers delivering at the University of Virginia Health System (n=8); consistency across the two sample populations was most evident among IgG1 glycan motifs, among which relationships between glycan motifs and maternal characteristics tended to trend in the same direction with a few correlations achieving statistical significance (p<0.05) (**Supplemental Figure S3**). These findings suggest maternal psychosocial and/or metabolic stress perturb the landscape of IgG glycosylation, which in turn disrupts placental transfer.

### Cell type-specific FcRn and FcγR expression signatures are consistent across gestation

While previous work provides evidence for both placental FcγRs and FcRn as determinants of antibody transfer efficiency and selectivity as a function of IgG Fc characteristics^5,15,26^, their cell type-specificity, colocalization, gestational expression trajectories, and relationship to Fc-specific IgG transfer have not been defined. To address these critical gaps, we performed mIHC staining of Fc receptors in placental tissue followed by quantitative image analysis in both cohorts (**Figure 3A**). Tissue sections from Cohort 1 were stained with panel 1, which contained cell type markers for STBs, ECs, and Hofbauer cells (HBCs), along with three Fc receptors of interest (**Figure 3A, Supplemental Figure S4**). Tissue sections from Cohort 2 were stained with panel 2, which was identical to panel 1 except for the inclusion of an anti-human IgG antibody in place of CD163 to analyze *in situ* colocalization between Fc receptors and IgG. Notably, although Cohorts 1 and 2 contained some specimens derived from pathological preterm births, the implication of preterm delivery did not appear to alter Fc receptor expression profiles (**Supplemental Figure S5**).

**Figure 3.**
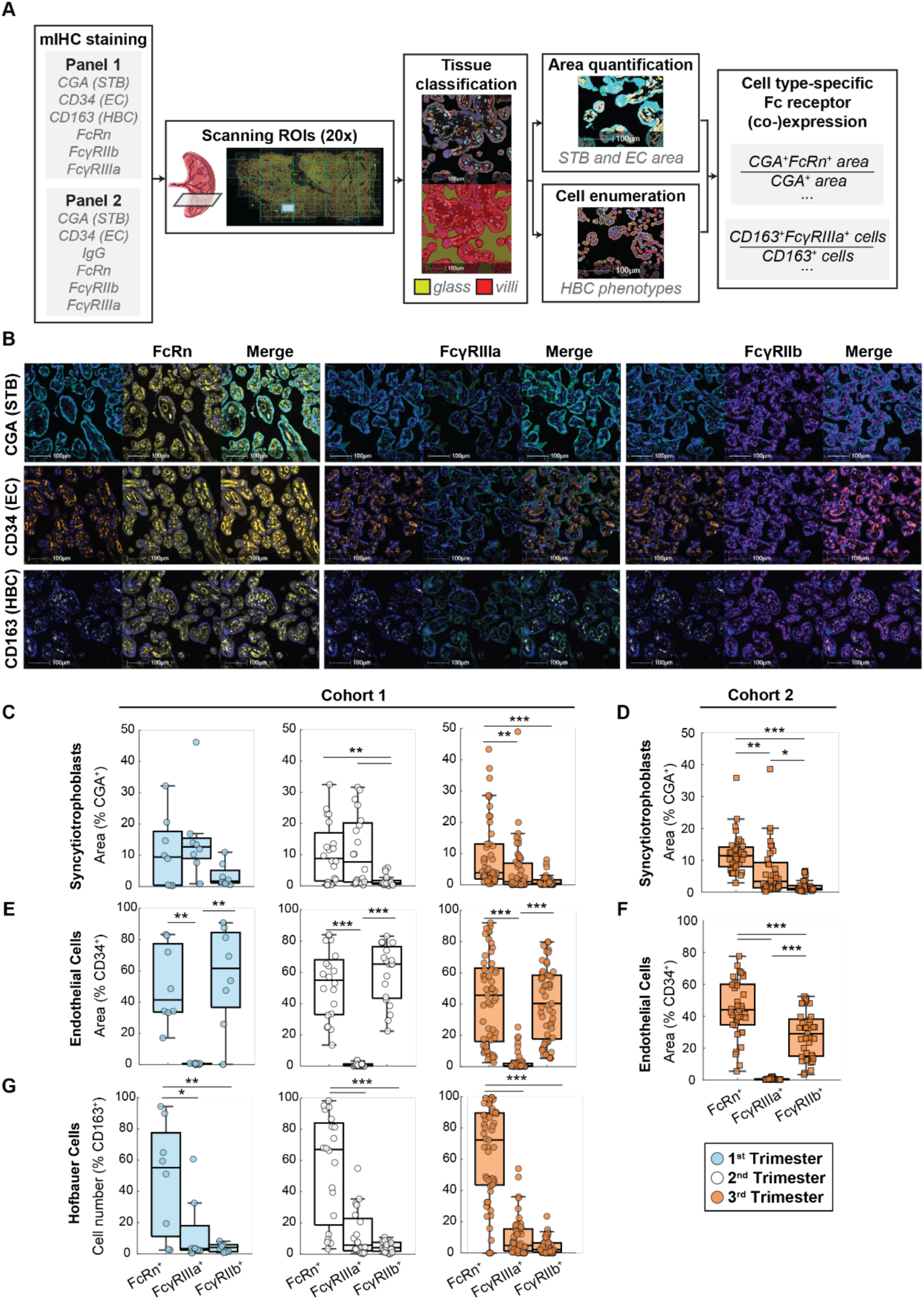
Trans-gestational profiling of placental Fc receptors by mIHC. (A) Quantitative image analysis pipeline. (B) Representative images exhibit colocalization patterns of each cell type marker with each Fc receptor in Panel 1. Scale bars, 100 μm. Stains are colored as follows, from left to right in order of appearance: CGA, cyan; FcRn, yellow; FcγRIIIa, green; FcγRIIb, magenta; CD34, orange; CD163, white. All images are shown overlaid with DAPI stain in blue. (C-G) FcRn, FcγRIIIa, and FcγRIIb expression quantified in situ on placental STBs (CGA^+^) in Cohort 1 (n=81) (C) and Cohort 2 (n=37) (D), on ECs (CD34^+^) in Cohort 1 (E) and Cohort 2 (F), and on HBCs (CD163^+^) in Cohort 1 (G). (***p<0.001, **p<0.01, *p<0.05, ANOVA with Tukey’s test for multiple comparisons). mIHC, multiplex immunohistochemistry. STB, syncytiotrophoblast. EC, endothelial cell. HBC, Hofbauer cell.

We quantified the colocalization of Fc receptors with cell type markers for ECs (CD34^+^), STBs (chorionic gonadotropin alpha, CGA^+^), and HBCs (CD163^+^)^27^ (**Figure 3B**). In both cohorts, STBs expressed primarily FcRn and FcγRIIIa, and <5% of CGA^+^ area colocalized with FcγRIIb across all trimesters (**Figure 3C-D**)^5,10,28,29^. In contrast, ECs expressed primarily FcRn and FcγRIIb as previously reported^28,30,31^, with some (<10% on average) FcγRIIIa expressed in samples from third trimester, especially in Cohort 1 (**Figure 3E-F**). On average, FcRn^+^ HBCs accounted for >50% of HBCs across all trimesters in Cohort 1 and were consistently more common than FcγRIIIa^+^ or FcγRIIb^+^ HBCs (**Figure 3G**). These data indicate that while FcRn is prominently expressed across STBs, ECs, and HBCs throughout gestation, FcγRIIb is restricted to ECs and FcγRIIIa is specific to STBs.

### Mechanistic insights derived from FcγR and FcRn colocalization analysis *in situ*

While several studies have hypothesized roles for FcγRIIb and FcγRIIIa in placental antibody transfer, the mechanisms by which these receptors regulate transfer and their potential interactions with FcRn have not been resolved. To address possible FcγR-FcRn co-regulation of IgG transfer, we quantified cell type-specific colocalization of FcγRs and FcRn in both cohorts. Interestingly, the majority of CGA^+^ and CD34^+^ area was classified as negative for both FcRn and FcγRIIIa (STBs) or FcγRIIb (ECs) (**Figure 4A-D**). Fc receptor expression is therefore not ubiquitous across the tissue but displays specific expression in areas which are putative antibody transfer hot spots.

**Figure 4.**
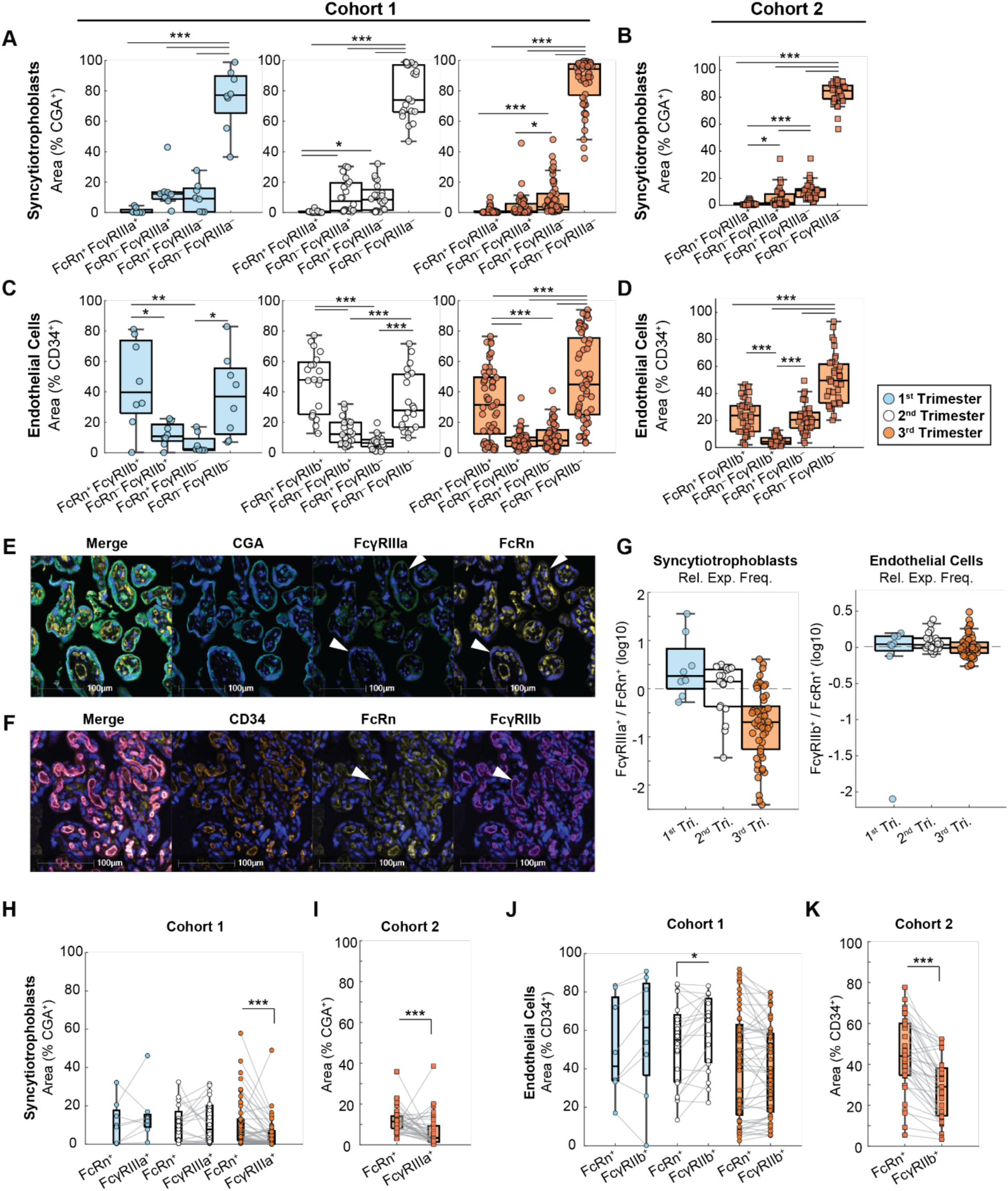
Spatial colocalization of FcRn and FcγRs in human placenta. (A-B) Area-based quantification of FcRn and FcγRIIIa colocalization with CGA in Cohort 1 (n=81) (A) and Cohort 2 (n=37) (B). (C-D) Area-based quantification of FcRn and FcγRIIb colocalization with CD34 in Cohort 1 (C) and Cohort 2 (D). (***p<0.001, **p<0.01, *p<0.05, ANOVA with Tukey’s test for multiple comparisons). (E) Representative image demonstrates mutually exclusive expression of FcRn and FcγRIIIa on CGA^+^ trophoblasts (white arrowheads). Stains are colored as follows, from left to right: CGA, cyan; FcγRIIIa, green; FcRn, yellow. All images are shown overlaid with DAPI stain in blue. Scale bars, 100 μm. (F) Representative image demonstrates abundant colocalization of FcRn and FcγRIIb on CD34^+^ vessels (white arrowheads). Stains are colored as follows, from left to right: CD34, orange; FcRn, yellow; FcγRIIb, magenta. All images are shown overlaid with DAPI stain in blue. Scale bars, 100 μm. (G) The log-transformed ratio of FcγRIIIa^+^ / FcRn^+^ CGA^+^ area (left) and FcγRIIb^+^ / FcRn^+^ CD34^+^ area (right) in Cohort 1. Dashed line at zero corresponds to a ratio of 1. (H-I) Paired analysis of CGA^+^ FcRn and FcγRIIIa colocalization in Cohort 1 (H) and Cohort 2 (I). (J-K) Paired analysis of CD34^+^ FcRn and FcγRIIb colocalization in Cohort 1 (J) and Cohort 2 (K). (***p<0.001, **p<0.01, *p<0.05, Wilcoxon signed rank test).

Although protein colocalization does not necessitate converging pathways of IgG transfer, we reasoned that the lack of colocalization between FcγRs and FcRn is consistent with an FcRn-independent, FcγR-mediated transfer mechanism. Across all three trimesters and in both cohorts, FcRn^+^FcγRIIIa^+^ colocalization was detected in <5% of CGA^+^ area (**Figure 4A-B**). Rather, the majority of FcγRIIIa^+^ area in STBs was not colocalized with FcRn, and vice versa. On the contrary, approximately 20-40% of CD34^+^ area colocalized with FcRn and FcγRIIb simultaneously (**Figure 4C-D**). In Cohort 2, FcRn was occasionally expressed without FcγRIIb, whereas FcγRIIb was rarely colocalized with CD34 in the absence of FcRn. In HBCs, the most common pair of Fc receptors to colocalize were FcRn and FcγRIIIa, yet Fc receptor colocalization occurred in <10% of HBCs on average (**Supplemental Figure S6**). Taken together, these data highlight the potential for both FcRn-dependent and -independent mechanisms of FcγRIIb-mediated transfer, while the lack of colocalization between FcRn and FcγRIIIa substantiates a possible FcγRIIIa-driven mechanism of IgG transcytosis beyond the canonical FcRn-mediated pathway (**Figure 4E-F**).

We have previously demonstrated that the relative expression of FcRn and FcγRIIb in placental ECs determines the propensity for preferential transfer of IgG1, IgG3, and IgG4 using a mathematical model^26^. Thus, we compared the expression frequency of FcγRIIIa and FcγRIIb relative to FcRn in STBs and ECs, respectively (**Figure 4G**). In Cohort 1, the frequency of CGA^+^ trophoblasts’ colocalization with FcγRIIIa relative to FcRn colocalization frequency decreased across trimesters. By the third trimester, FcγRIIIa expression frequency was significantly less than FcRn frequency (FcγRIIIa/FcRn=0.44, *p*<0.001) (**Figure 4G-H**). A similar pattern was observed in the third trimester samples from Cohort 2 (FcγRIIIa/FcRn=0.54, *p*<0.001) (**Figure 4I**). On the other hand, FcγRIIb expression was slightly more common than FcRn in second trimester patient tissues in Cohort 1 (FcγRIIb/FcRn=1.14, *p*<0.05), but the two receptors were expressed by a similar fraction of ECs in the first and third trimester (**Figure 4G,J**). In Cohort 2, ECs expressed FcRn more prevalently than FcγRIIb (FcγRIIb/FcRn=0.63, *p*<0.001) (**Figure 4K**). In sum, during second trimester a similar fraction of STBs and ECs express FcγRIIIa and FcγRIIb compared to FcRn, but both cell types exhibit an FcRn-dominant expression profile approaching term.

### FcγRs are enriched in first and second trimester human placenta

Previous studies have shown that IgG in preterm and term newborns is of similar functional quality, suggesting antibodies are transferred at non-constant rates across pregnancy resulting in early, preferential transfer of functional antibodies^15^. We hypothesized that variable Fc receptor expression in key placental cell types across gestation temporally regulates maternal-fetal IgG transfer. To define Fc receptor expression dynamics, we plotted the average receptor expression frequency per cell type in each specimen from Cohort 1 as a function of gestational age (**Supplemental Figure S7**). Most receptors—especially those expressed by STBs and ECs—exhibited non-monotonic trajectories across gestation. Notably, 3^rd^-order polynomial regression models for FcRn^+^ and FcγRIIIa^+^ trophoblasts and FcRn^+^ and FcγRIIb^+^ ECs predicted peak expression frequency at approximately 20 weeks’ gestational age, consistent with their contribution to delivering functional, FcγR-binding antibodies to premature neonates.

To elucidate patterns of Fc receptor expression variation across gestation, we performed orthogonalized PLSDA modeling to predict each specimens’ gestational age from the corresponding receptor expression profile in Cohort 1 (**Figure 5A-D**) and validated these findings in Cohort 2 (**Figure 5E-F**). Given the aforementioned nonlinear expression dynamics, we trained a model to predict each sample’s trimester rather than absolute gestational age to allow detection of non-monotonic changes. The resulting PLSDA model discriminated trimesters with a mean cross-validation accuracy of 73% and was statistically significant by permutation test (*p*<0.001) (**Figure 5A**). Examination of loadings on latent variable 1 (LV1), which captured the variance with gestational age, along with VIP score analysis revealed gestational age-associated features that discriminated across trimesters (**Figure 5B-C**). These features included FcγRIIIa and FcγRIIb expressed by STBs, which were expressed most prominently in first trimester, and FcRn expressed by HBCs which exhibited a subtle increase across trimesters. Univariate analysis revealed a subset of these features, including FcγRIIIa^+^ STBs and FcγRIIb^+^ STBs and ECs, that were significantly enriched in samples from the first and/or second trimesters compared to third trimester (**Figure 5D**). In contrast, none of the HBC-linked features were significantly variable across trimesters, as anticipated by their poor univariate regression scores (R^2^<0.1) (**Supplemental Figure S7**). Nevertheless, these tightly coordinated changes across gestation indicate temporal, tissue-level regulation of Fc receptor expression.

**Figure 5.**
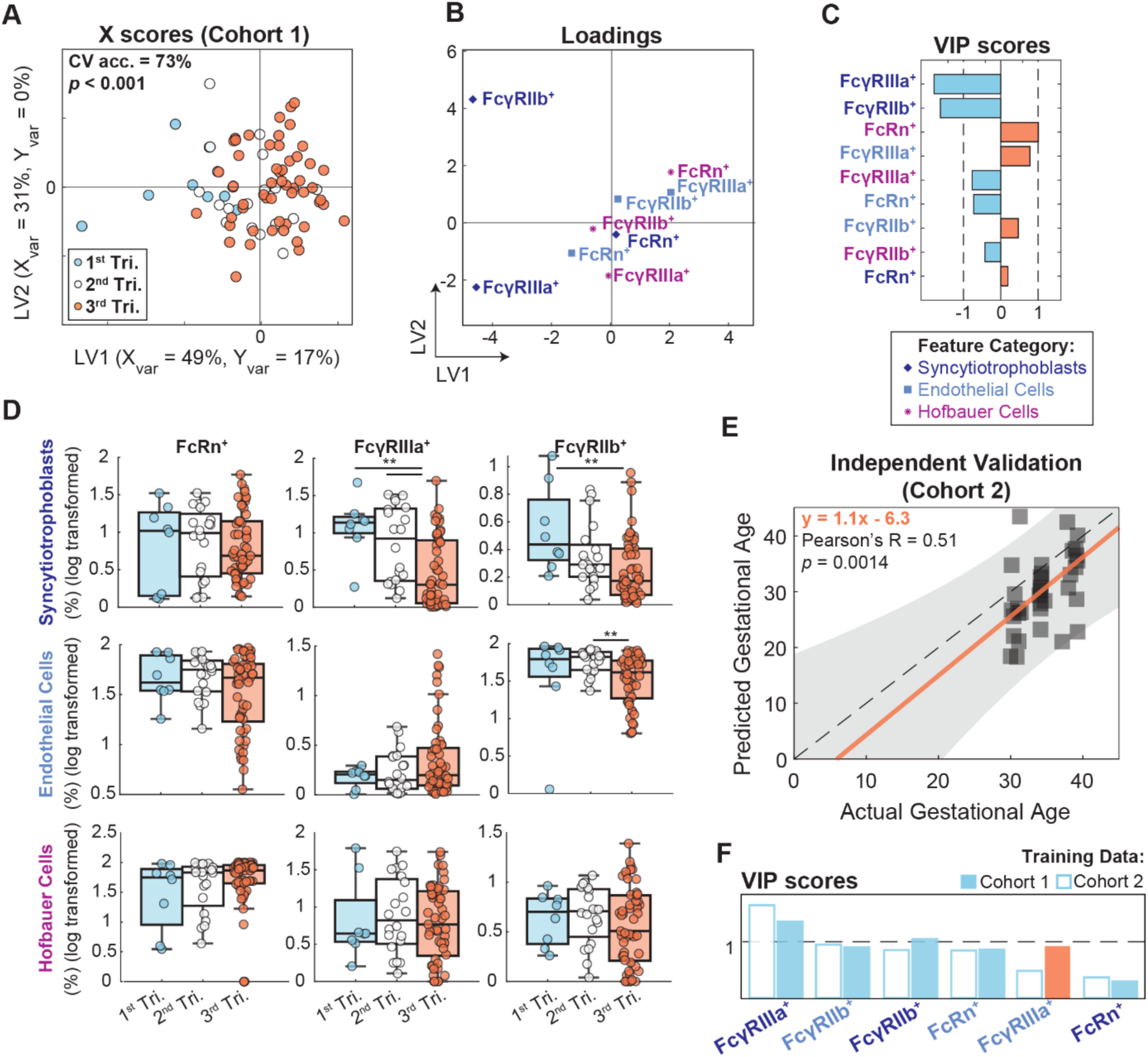
Temporal trajectories of Fc receptors in human placenta across gestation. (A) X scores plot from a 3-class PLSDA model predicting specimen trimester from placental Fc receptor expression frequencies in Cohort 1 (n=81). The variance in X and Y captured by each latent variable (LV1 and LV2) is denoted on the axis labels. Significance was determined by a two-sided t-test comparing the true model accuracy scores over 1,000 independent rounds of 5-fold cross-validation against 1,000 models trained on data with randomly permutated class labels. (B) Feature loadings on LV1 and LV2 are colored and marked with a symbol corresponding to their cell type specificity. Indigo diamonds, STBs. Light blue squares, ECs. Fuchsia asterisks, HBCs. (C) VIP scores bar plot displays the relative importance of each feature to model performance. VIP > 1 denotes features with greater than average influence on the model. VIP scores are artificially oriented and colored according to their loadings on LV1. (D) Univariate box plots comparing cell-specific Fc receptor expression across trimesters. (*p<0.05, **p<0.01, ***p<0.001, Kruskal-Wallis with Dunn’s test for multiple comparisons). (E) An OPLSR model was trained on the intersect of features in cohorts 1 and 2 quantifying Fc receptor expression frequency on STBs and ECs (6 features). The scatter plot shows predicted vs. actual gestational age in Cohort 2 (n=37) using the OPLSR model trained on data from Cohort 1. Each point represents one patient tissue. The dotted line represents y=x. The orange line is a linear regression shown with 95% confidence intervals. (F) VIP scores compared across two independent OPLSR models trained on data from Cohort 1 (filled bars, corresponding to the model used for prediction in (E)) and Cohort 2 (open bars). The feature labels are colored according to their cell type. Bars are ordered based on VIP score rankings in the Cohort 2 model and colored according to their loadings on LV1 (blue = negative association with gestational age, orange = positive association with gestational age). PLSDA, partial least squares discriminant analysis. CV acc., cross validation accuracy. STB, syncytiotrophoblast. EC, endothelial cell. HBC, Hofbauer cell. VIP, variable importance in projection. OPLSR, orthogonalized partial least squares regression.

To validate these inferred dynamic trends, we trained an orthogonalized partial least squares regression (OPLSR) model on data from Cohort 1 using only the shared features from Cohorts 1 and 2. The OPLSR model displayed strong predictive power and was statistically significant by permutation testing (Q^2^=0.41, *p*<0.001) (**Supplemental Figure S8**). We performed independent model validation by predicting gestational age in Cohort 2; the model validation was robust, exhibiting a significant linear correlation between predicted and actual gestational age (Pearson’s R=0.51, *p*=0.0014, regression slope=1.1) (**Figure 5E**). Given that Cohort 2 spanned only the third trimester, this signified a dynamic shift in placental Fc receptor expression evident even in the narrow window of third trimester when a substantial degree of antibody transfer occurs. To validate the dynamic trajectories of individual features across cohorts, we trained a second OPLSR model on data from Cohort 2 and compared the VIP scores to those from the Cohort 1-trained model (**Figure 5F**, **Supplemental Figure S8**). In both models, all features had positive loadings indicating an increase with gestational age, apart from FcγRIIIa^+^ ECs which increased in Cohort 1 only (**Figure 5F**, orange solid bar). The top three features in both models were FcγRIIIa^+^ STBs, FcγRIIb^+^ STBs, and FcγRIIb^+^ ECs (VIP>1), suggesting both barrier cell types downregulate Fc receptor expression as gestation approaches term. In sum, these data point to coordinated, cell-specific Fc receptor expression dynamics across gestation consistent across both sample populations.

### Placental FcγRs explain variation in IgG subclass- and glycan-specific transfer

As maternal subclass-specific glycosylation affected placental transfer efficiency and was apparently perturbed by clinical features, we hypothesized that placental Fc receptors may similarly be impacted by these key covariates. However, we did not detect significant variation in Fc receptor expression across the patient clusters identified in **Figure 1** in either cohort (**Supplemental Figure S9**), indicating that maternal IgG Fc features are more directly linked to variation across clinical profiles. Regardless, we hypothesized that Fc receptors in placenta serve as the mechanistic link between maternal Fc variability and placental transfer efficiency by selectively transferring high-affinity antibodies. To evaluate relationships between Fc receptor expression profiles and Fc-specific antibody transfer, we performed partial correlation network (PCN) analyses. Because we observed both Fc receptor expression and cord IgG subclass levels vary with gestational age (**Figure 5**, **Supplemental Figure S10**), we controlled for gestational age and other clinical covariates which impacted antibody transfer (maternal age, gravida, pregravid BMI, and EPDS scores). Importantly, controlling for these covariates resulted in stronger correlations and changed the directionality of some correlations, likely due to inverse relationships between gestational age and Fc receptor expression (**Supplemental Figure S11**).

We first assessed correlations between cell type-specific Fc receptor expression frequencies and subclass-specific transfer (**Figure 6A**). In Cohort 1, we found FcRn^+^ STBs positively correlated with both cord concentrations and transfer ratios of all four IgG subclasses, consistent with the established role of FcRn in IgG transcytosis across STBs^32^ (**Supplemental Figure S11**). Likewise, FcRn^+^ and FcγRIIb^+^ ECs as well as FcγRIIIa^+^ STBs positively correlated with subclass transfer ratios and cord concentrations. The strongest correlations were FcRn^+^ and FcγRIIb^+^ ECs with cord IgG4 concentrations, FcRn^+^ STBs with cord IgG3 and IgG2 concentrations, and FcγRIIb^+^ HBCs with cord IgG1 concentrations (*p*<0.1) (**Figure 6A**). Interestingly, FcγRIIIa^+^ HBCs were inversely correlated with transfer efficiency of all subclasses, possibly indicating a role in phagocytosis^33^ (**Supplemental Figure S11B**, not significant). In Cohort 2, Fc receptor expression frequencies exhibited similar strong correlations with IgG subclass transfer ratios, but not absolute cord concentrations. FcRn^+^ STBs, FcRn^+^ ECs, and FcγRIIb^+^ ECs positively correlated with IgG2 cord:maternal ratio (p<0.1), and FcRn^+^ ECs was also correlated with IgG1 and IgG3 ratios (**Figure 6A**). Taken together, these relationships indicate subclass-specific effects of both FcRn and FcγRs on antibody transfer efficiency.

**Figure 6.**
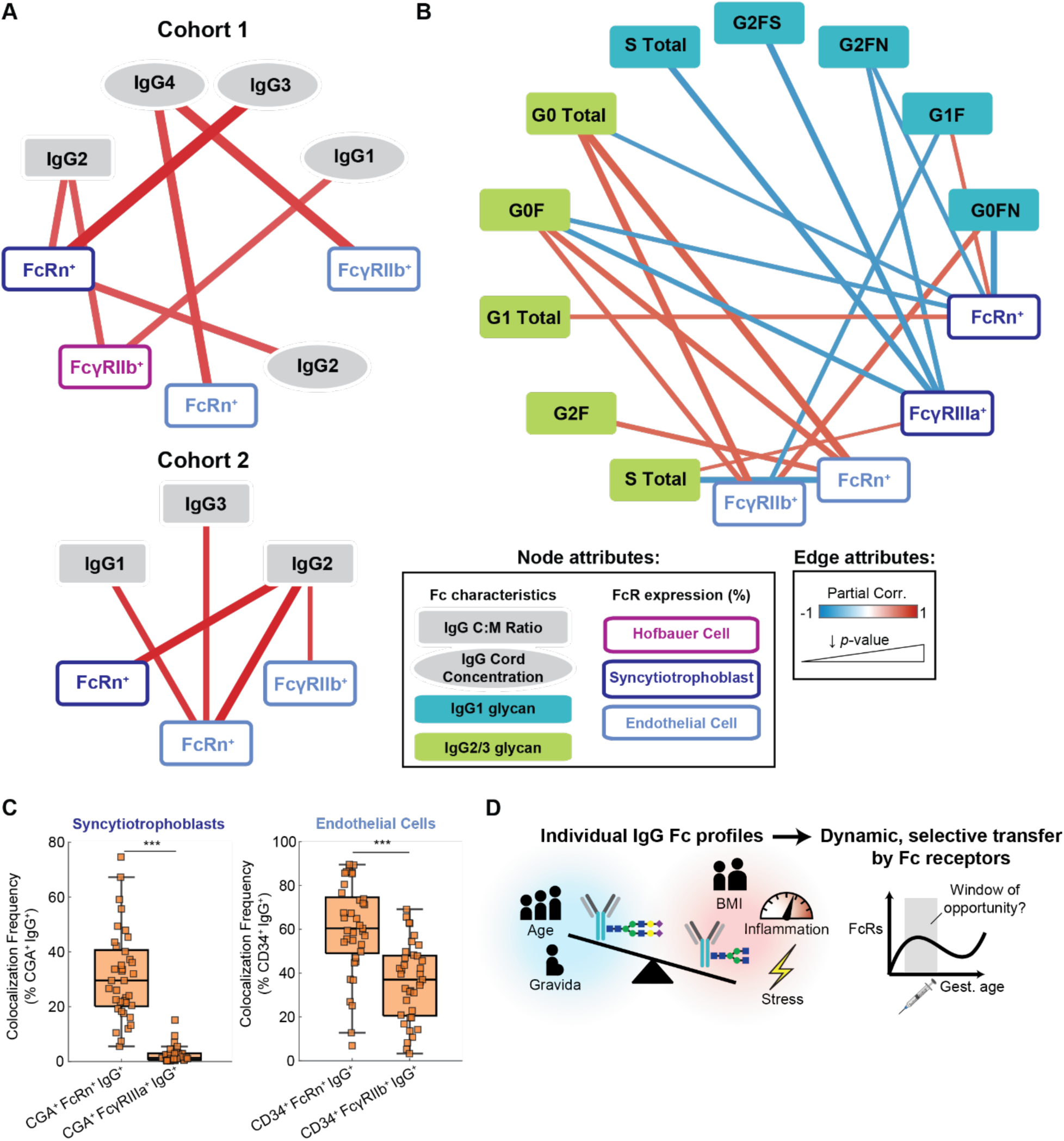
Partial correlation network analysis reveals relationships between placental Fc receptor expression and maternal Fc-specific antibody transfer in matched samples. (A) Partial correlation network between IgG subclass C:M transfer ratios (gray rectangles), absolute cord concentrations (gray ellipses), and cell type-specific Fc receptor expression frequencies in Cohort 1 (top, n=16) and Cohort 2 (bottom, n=10). Fc receptor nodes are colored according to cell type. The legend describes encoding of node and edge attributes for both networks. Edge color corresponds to the partial correlation coefficient (red, positive correlation; blue, negative correlation) and edge width is inversely proportional to the correlation p-value. Only correlations with (p<0.1) are shown. (B) Partial correlation network shows the relationships between Fc receptor expression frequencies and glycan- and subclass-specific transfer ratios in matched specimens from Cohort 2 (n=8). Nodes are colored according to IgG subclass or Fc receptor cell type. The legend from (A) applies. (C) Cell-specific colocalization frequency of each receptor with IgG quantified in syncytiotrophoblasts (left) or endothelial cells (right) in mIHC images from Cohort 2. (***p<0.001, Wilcoxon test). C:M ratio, cord:maternal ratio. (D) Conceptual model summarizes the key insights from this paper. Maternal clinical covariates determine the Fc subclass and glycan profile. Placental Fc receptors selectively transfer IgG based on Fc characteristics, and the dynamic regulation of this process across gestation indicates a vaccination window of opportunity during gestation.

We next performed PCN analysis for patients with matched placental Fc receptor expression profiles and IgG subclass glycans (n=8) (**Figure 6B**). We found that FcγRIIb and FcRn expression frequency in ECs was typically positively correlated with glycan species’ transfer efficiency, including those which displayed inefficient antibody transfer (e.g., IgG1 G0FN, IgG2/3 G0F) and some which displayed efficient transfer (IgG2/3 G2F) (**Supplemental Figure S2**). This is consistent with a role of EC Fc receptors as rate-determining factors in placental transfer; when Fc receptors are available in excess of maternal IgG, the likelihood of transferring low-affinity antibodies increases^26^. Meanwhile, we observed primarily negative correlations between FcγRIIIa^+^ STBs and various glycan species, even those displaying efficient transfer (e.g., IgG1 G2FS and G2FN). Similar relationships were observed for FcRn^+^ STBs, with additional positive correlations detected with monogalactosylated IgG1 (G1F) and IgG2/3.

To glean additional insights into the relative contribution of each Fc receptor to IgG transfer across STBs and ECs, we quantified the cell-specific colocalization of each receptor with IgG *in situ* in Cohort 2 (**Figure 6C**). Around 30% of IgG colocalized with CGA^+^ tissue (STBs) and 60% of IgG colocalized with CD34^+^ tissue (ECs) also colocalized with FcRn, which suggests a strong contribution of FcRn to IgG transfer in both cell types. An additional ∼30% of IgG colocalized with FcγRIIb in ECs, implicating FcγRIIb as a supplementary regulator of IgG transcytosis across this barrier. Consistent with the PCN analysis, these data support a key role for FcRn driving IgG transfer across both STBs and ECs independent of IgG subclass, whereas FcγRIIb and FcγRIIIa may enhance transfer of low-abundance, high-affinity subclasses and glycoforms. Therefore, a combination of placental FcγR expression and maternal IgG Fc characteristics (subclass, glycosylation) jointly determine antibody transfer efficiency and may underscore interpersonal heterogeneity in transfer dynamics across gestation (**Figure 6D**).

## Discussion

IgG transplacental transfer plays a critical role in protecting newborns against infectious diseases^34–36^. The mechanisms which give rise to heterogeneity in the quantity, antigenic repertoire, and functionality of antibodies passed to the newborn are incompletely understood^8^, but include a combination of maternal and placental factors. Previous studies have provided evidence for glycan-specific antibody transfer ^4,5^, while others have demonstrated abundant FcγR expression in placental villi^4,5,11^ and postulated the role of these receptors in transfer^37^. Further, recent evidence has demonstrated a requirement for an additional Fc receptor beyond FcRn regulating placental IgG transfer in non-human primates, but not mice, suggesting alternative receptor(s) may likewise play a key role in placental antibody transfer in humans^38,39^. In this study, we holistically dissected regulators of maternal-fetal antibody transfer by leveraging two human cohorts spanning a broad range of gestational ages with matched maternal, cord, and placental tissues and with multimodal measurements including clinical covariates, IgG subclass-specific Fc glycosylation, and *in situ* Fc receptor expression. This analysis offered unprecedented insight into multifactorial regulatory networks involving maternal and placental factors which explain population level heterogeneity in placental antibody transfer^3^. We found that readily available clinical data alone explained inter-personal variability in subclass-specific transfer efficiency, revealing three patient profiles stratified by their clinical features and IgG transfer signatures. There were robust correlations between key clinical covariates (e.g., age, pregravid BMI, CRP, EPDS score) and subclass-specific glycosylation, suggesting maternal social and metabolic stressors alter Fc glycosylation which in turn affects placental transfer efficiency.

Importantly, placental Fc receptors were not as strongly associated with the clinical covariates. However, FcγRs might underscore the mechanism by which IgG glycans influence transfer due to varying affinities of FcγRs for glycosylated IgG^40,41^, and subclass- and glycan-specific transfer rates were correlated with FcRn and FcγR expression frequencies in matched placental tissues. This highlights a role for Fc receptors as rate-determining factors in selective antibody transfer. We found these receptors displayed cell type-specific expression patterns which were conserved across gestation and were consistent in both cohorts. Comparing Fc receptor expression across tissues from different gestational time points revealed a shift from an FcγRIIb- and FcγRIIIa-rich landscape during first trimester to a predominantly FcRn-rich landscape during third trimester. These trends reflect potential temporal regulation of placental sieving activity, consistent with previous findings that functionally active antibodies are present in similar proportions in preterm and term babies, albeit at lower absolute titers^15^.

IgG Fc glycosylation differences between maternal and cord plasma have been investigated over the past several decades in multiple cohorts and with varying glycan analysis techniques. Here, we found that terminal sialylation and digalactosylation across all IgG subclasses were enriched in cord compared to their matched maternal levels. Pregnancy is known to be associated with higher levels of galactosylation and sialylation^42–44^, which is advantageous given the selective transfer of these specific IgG antibodies. The preferential transfer of galactoslated IgG Fc glycans was previously observed by us and others^5,6,16,39,45–47^. Sialylated IgG Fc glycan differences between maternal and cord plasma IgG were also found to be significant^6^ or trending higher in previous studies^5,39^. One study analyzed ten pairs of fetal and maternal IgG samples revealing largely comparable Fc glycosylation for all the analyzed subclasses^48^, speculating that the difference between their study and earlier ones might stem from the differences in total (Fc plus Fab) vs. Fc glycosylation. IgG Fab glycans were previously shown to hinder FcRn-mediated placental transfer^49^. However, a low percentage of IgG (approximately 20% of all immunoglobulins) is Fab-glycosylated in healthy people, making it an important consideration only in malignant or autoimmune conditions harboring a higher Fab glycosylation^50^. While some of these studies that provided evidence for glycan-mediated transfer analyzed total IgG glycosylation^16,46,51,524–6^, more recent studies focused on Fc glycans^4–6^. Our methods specifically measured Fc glycans and confirm these latter studies. The present study quantifies IgG subclass-specific glycans using LC-MS glycoproteomics analysis, a highly sensitive method, to resolve subclass-specific Fc glycoforms (See Methods). Jansen et al found lower per antenna and per galactose sialylation of IgG (total Fc and Fab sialylation) in cord compared to maternal levels^45^.

The mechanisms underpinning temporal regulation of placental antibody transfer in humans have not been described, in part due to restrictions to studying this process in human cohorts and inter-species anatomical differences which limit the translational relevance of rodent and primate models^53^. We observed expression frequency of most placental Fc receptors peaked during second trimester, potentially defining a vaccination window of opportunity to maximize antibody transfer to the fetus by capitalizing on early, efficient transfer of functional antibodies^39^. Consistent with this model, prenatal SARS-CoV-2 mRNA vaccination timing has been found to affect the quality and durability of anti-RBD and anti-spike IgG in newborns, revealing a vaccination “sweet spot” to maximize not only the titer at birth, but also functionality and protective benefit for newborns throughout the vulnerable period of early infancy^8^. Notably, other studies report effects of SARS-CoV-2 vaccination timing on placental transfer efficiency, with earlier vaccination resulting in more efficient vaccine-induced antibody transfer compared to third trimester vaccination^54,55^. Similar effects were also reported for prenatal Tdap vaccines^56,57^. However, one such report^55^ showed decreased functional scores and Fc receptor binding capacity following second trimester vaccination, perhaps due to an anti-inflammatory maternal immune phenotype during second trimester which is necessary to support fetal growth^58^. Therefore, there exists a window of opportunity in mid-gestation to transfer functional, FcγR-binding antibodies following maternal vaccination, but this must be balanced with the natural variation in maternal vaccine responses across gestation.

Together, these data demonstrate a complex interplay between environmental, immune, and placental features driving population heterogeneity in antibody transfer. We observed similar effects of digalactosylation on placental sieving as reported^4,5^; still, variation of IgG glycosylation profiles across diverse cohorts is well-documented^22,48,59,60^ These insights could impact clinical practice by recommending lifestyle modifications and personalized vaccine approaches to shift the IgG glycome toward a profile conducive to placental transfer. For instance, physical exercise, dietary modifications, and weight loss have been shown to increase IgG digalactosylation and sialylation, both of which are associated with enhanced antibody transfer^61–63^. Alternatively, vaccines can be engineered to induce different IgG glycosylation profiles by modulating design parameters including adjuvant, dosage, and target antigen^64–67^. This presents an opportunity to formulate pregnancy-specific vaccines that induce IgG responses conducive to efficient placental transfer. As we await the validation of reliable and accessible biomarkers of placental Fc receptor expression and/or transfer efficiency, the translational potential of predicting IgG transfer efficiency from readily available maternal clinical data should be emphasized.

Our study design was robust due to parallel analysis of two independent cohorts, high-resolution LC-MS glycan analysis, high-throughput imaging techniques, and analysis of matched plasma and placental tissue. Regardless, there are limitations to consider. First, our mIHC panels were limited to six protein targets plus DAPI. This prevented us from conducting an unbiased investigation of all FcγR types and isoforms and disentangling expression variation across different trophoblast lineages and HBC phenotypes. Second, our cohort comprised primarily healthy tissues, and we did not investigate the effect of perturbations such as infections or prenatal immunizations on Fc receptor expression and antibody transfer efficiency, an important consideration given findings from others that placental Fc receptor expression is perturbed in viral infections such as SARS-CoV-2^10^. Third, our cross-sectional approach to mapping the temporal trajectories of Fc receptor expression across gestation assumes that tissues from different patients and time points reflect longitudinal placental development, which may not hold especially when considering specimens from pathological preterm births as a proxy for healthy second and early third trimester tissues. Finally, our study is observational in nature, and we performed only associative statistical modeling on these data; future mechanistic experiments are needed to fully resolve the role of FcγRIIb and FcγRIIIa and their potential interplay with FcRn to modulate subclass- and glycan-specific antibody transfer. Still, this is one of few studies to report associations between placental Fc receptor expression and IgG transfer efficiency, substantiating a critical role for these receptors as drivers of Fc-specific antibody transfer. Future work can expand upon these findings by integrating the heterogeneity and expression dynamics into mechanistic models of placental antibody transfer to refine predictions of precisely timed, precision prenatal immunization approaches.

Ultimately, our study reveals multimodal regulatory networks governing subclass- and glycan-specific antibody transfer and its dynamic regulation across human pregnancy. These insights provide a basis for the rational design of maternal vaccines with enhanced binding to FcγRs for optimized placental transfer. Moreover, our data provide a rationale for immunization in early gestation where appropriate and safe for the mother and fetus to enhance functional antibody transfer. Our study constitutes an important step toward disentangling placental IgG transfer variation across the population and the eventual implementation of personalized prenatal immunizations that offer optimal protection to newborns against existing and emerging infectious diseases.

## Methods

### Patient population

A prospective cohort of pregnant persons was enrolled at the University of California, Los Angeles (UCLA) Perinatal Biorepository (ClinicalTrial.gov: NCT05035160) (**Table 1**). The study was approved by the Institutional Review Board (UCLA IRB# 21-001018). Eligible participants were pregnant people willing to give informed consent. After the patient was identified, participants underwent screening and eligible participants were contacted by a study coordinator by phone or email. Informed consent was obtained, and Health Insurance Portability and Accountability Act (HIPAA) release of medical records was obtained electronically using DocuSign. Demographic and clinical data were extracted from electronic health record and maintained in a de-identified database. Samples from the 1^st^ and 2^nd^ trimester were obtained from patients who either underwent an elective termination or spontaneous abortion. A subset of these patients did not consent to medical record release and thus the cause of pregnancy termination was unavailable. A suspected congenital abnormality was reported for eight of the 1^st^ and 2^nd^ trimester cases; Fc receptor expression was not found to vary with suspected fetal anomaly.

A second prospective cohort was enrolled at the University of Virginia (UVA) and approved by the Institutional Review Board (UVA IRB #210420) (**Table 2**). Eligible participants were pregnant people willing to give informed consent. After a patient was identified, the patient underwent screening and eligible participants were contacted by a study coordinator using MyChart. Informed consent was obtained. Demographic and clinical data were extracted from electronic health record and maintained in a de-identified database. Additional formalin-fixed, paraffin embedded full-thickness placental tissue sections from grossly normal appearing tissues were obtained retrospectively through the Biorepository and Tissue Research Facility at UVA (UVA IRB #210416).

### Sample preparation

Placental specimens were processed as soon as possible and generally within 4 hours of delivery. Placental biopsies were extracted using standard operating procedures, washed 3 times with phosphate-buffered saline (PBS), and fixed with 10% paraformaldehyde overnight (12-16 hours). Full-thickness sections of the grossly normal-appearing placenta from the chorionic plate to the basal plate were obtained. The sections were embedded in paraffin and stored at room temperature. For each prospectively collected placental sample from both cohorts, maternal blood was collected directly on the birth admission concomitant with placement on the peripheral IV. Birth hospitalization is defined as the admission with the birth. Umbilical cord blood was drawn immediately following delivery of the placenta. Maternal and umbilical cord plasma was isolated by centrifugation and stored in single-use aliquots at <–70°C until analysis.

### Multiplex immunohistochemistry

4 μm-thick formalin-fixed, paraffin-embedded (FFPE) sections of human placental tissue were prepared fresh within two weeks of staining. mIHC was performed according to the manufacturer’s protocol using the OPAL Multiplex Manual IHC kit and antigen retrieval (AR) buffers AR6 and AR9 (Akoya Biosciences, Marlborough, Massachusetts, USA) and DIVA Decloaker AR buffer (Biocare Medical, Pacheco, California, USA).

Cohort 1 was stained, scanned, and analyzed in 2025. The staining sequence for Panel 1 was as follows: AR9, FCGRT/FcRn α-chain (1:200, LSBio C408078, rabbit, polyclonal) Opal 570; AR6, CD163 (1:200, Abcam ab182422, rabbit, monoclonal) Opal 650; AR9, CGA (1:100, Atlas Antibodies HPA029698, rabbit, polyclonal) Opal 690; AR9, CD16A/FcγRIIIa (1:100, Abcam ab227665, rabbit, monoclonal) Opal 520; AR6, CD32B/FcγRIIb (1:100, Abcam ab45143, rabbit, monoclonal) Opal 540; DIVA, CD34 (1:200, DAKO M7165, mouse, monoclonal) Opal 620; AR6, spectral DAPI (Akoya Biosciences). Cohort 2 was stained, scanned, and analyzed in 2024. The staining sequence for Panel 2 was as follows: DIVA, FCGRT/FcRn α-chain (1:100, Abcam ab2228975, rabbit, monoclonal) Opal 570; AR6, IgG (1:10,000, Abcam ab109489, clone EPR4421, rabbit, monoclonal) Opal 650; AR9, CGA (1:100, Atlas Antibodies HPA029698, rabbit, polyclonal) Opal 690; AR9, CD16A/FcγRIIIa (1:100, Abcam ab227665, rabbit, monoclonal) Opal 520; AR6, CD32B/FcγRIIb (1:100, Abcam ab45143, rabbit, monoclonal) Opal 540; DIVA, CD34 (1:200, DAKO M7165, mouse, monoclonal) Opal 620; spectral DAPI (Akoya Biosciences).

Stained slides were mounted using prolonged diamond antifade (Life Technologies, Carlsbad, California, USA) and scanned using the PerkinElmer Vectra 3.0 system and Vectra software (Akoya Biosciences). Regions of interest (3 mm^2^) were identified using Phenochart software and tissue images were acquired at 20x magnification. Regions were selected to avoid images containing staining artifacts (e.g., bubbles, folded tissue). The images were spectrally unmixed using InForm software (Akoya Biosciences) based on single stain positive control tissue sections from human placenta, tonsil, and tumor-involved lymph node. Tissue stained with no primary antibody were used as negative controls.

After scanning, images were visually inspected to assess stain quality. 19 of 100 tissues from Cohort 1 displaying irregular and weak staining of canonical cell type markers in the villi (CD34, CGA, and CD163) were excluded from downstream analysis.

### Quantitative image analysis

Image analysis was performed in HALO software v.4.0 (Indica Labs). For both cohorts, a random forest tissue classifier algorithm was trained to delineate glass, villous tissue, and non-villous tissue (e.g., decidua, fetal membranes, parenchyma) based on staining patterns of CGA and CD34. Subsequent quantifications were performed only on villous tissue. Quantification algorithms were applied to whole images (3 mm^2^) and the mean expression across all images per patient were used in downstream analyses. The Area Quantification algorithm was used to quantify colocalization of CGA (STBs) or CD34 (ECs) with one or more Fc receptors, yielding cell-type specific receptor expression and co-expression frequency profiles in both cohorts. For example, the frequency of FcRn-positive trophoblasts was defined as:

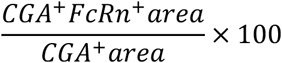

For Cohort 1, the HighPlex FL algorithm was employed to segment and enumerate CD163^+^ macrophages in the villous stroma (HBCs). Downstream analyses were performed on the cell density summary data, expressed as cells/mm^2^. The frequency of each receptor expression was determined as a percentage of HBCs expressing the receptor normalized to the total HBC density. For example, the frequency of FcRn-positive HBCs was defined as:

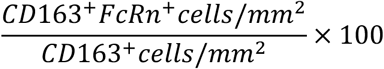

Area-based expression frequency and cell-based expression frequency (Cohort 1 only) were combined into a single data set for downstream analysis.

### Immunoglobulin isotyping assay

Immunoglobulin isotypes were quantified in a subset of maternal and umbilical cord plasma samples from both the cohorts using the Milliplex Map Human Isotyping Magnetic Bead Panel kit (HGAMMAG-301K, Millipore Sigma) per the manufacturer’s instructions. Plasma samples were diluted 1:15,000 in assay buffer. Samples were read on the Luminex MagPlex machine courtesy of UVA Flow Cytometry Core Facility. Absolute concentrations of IgG1, IgG2, IgG3, and IgG4 were determined based on 5-parameter logistic regression standard curves for each isotype.

### Plasma IgG glycoproteomics

Glycoproteomics analysis was performed at the University of Georgia Complex Carbohydrate Research Center (CCRC). Briefly, IgG subclass-specific glycans were quantified in matched maternal and umbilical cord plasma at the time of delivery from a subset of Cohort 2 (n=14 dyads) using liquid chromatography-mass spectrometry (LC-MS). A separate cohort of mothers (UVA IRB #19114) (n=8) with plasma collected at the time of delivery were obtained retrospectively and analyzed with the same method for validation.

#### Sample Preparation

Five microliter patient plasma (∼300 µg total plasma proteins) was drawn from each sample and diluted with 250 µL 50 mM ammonium bicarbonate (ABC). To the sample solution was added 200 mM D,L-dithiothreitol (DTT) to a final concentration of 5 mM. The mixture was incubated at 50°C for 45 min. The denatured sample was treated with 200 mM α-iodoacetamide (IAA) to final IAA concentration of 13 mM, and the sample was incubated at room temperature in dark for 45 min. Excess IAA was quenched by adding 200 mM DTT to a final concentration of 5 mM, followed by incubation at room temperature for 15 min. Approx. 30 µg plasma protein (28 µL) in ABC solution was treated with 3 µL 0.4 µg µL^-1^ sequencing grade Trypsin (1:25 enzyme:substrate ratio) and incubated at 37 °C overnight. Digested samples were diluted with 0.1% formic acid (FA) to 0.1 µg µL^-1^ (calculated from ∼300 µg total starting plasma proteins in each sample) and filtered through 0.2 µm filters.

#### LC-MS Parameters

LC-MS analysis was performed on Vanquish Neo – Orbitrap Ascend system using an Acclaim PepMax 100 C18 nanoViper column (75 µm × 15 cm, 2 µm). Peptides were separated in 1 h LC-MS/MS runs where Buffer B (80% Acetonitrile, ACN 0.1% FA) was ramped 4-25% in 0-40 min, flowrate 0.350 µL min^-1^ (Buffer A is 0.1% FA). The column was cleaned up by 99% Buffer B in 40-55 min and equilibrated to 4% Buffer B in 55-60 min. MS1 m/z was scanned in the range of 350-2000, RF lens 65%, and Orbitrap resolution 120k. Profile peaks were recorded. A data-dependent acquisition program was used for MS2. This aided glycopeptide retention time (RT) determination. Precursors were excluded for 30 s after being observed twice in 10 s. Precursor charges 1-6 with minimum intensity 5.0e4 were sent to steppedHCDpdsteppedHCD (HCD, higher collision energy dissociation; pd, product dependent) fragmentation^68–70^. The initial round of 20%/35%/50% normalized collision energy (NCE) HCD generated Y1 (Pep+HexNAc(1)) and Y1F (Pep+HexNAc(1)Fuc(1)) ions of IgG1, IgG2/3a, IgG3b/4 clusters, which triggered 25/30/45% NCE HCD to maximize peptide backbone b/y ions^71–73^. Approx. 0.1 µg peptides were injected for each sample. No blank injection is necessary between sample injections. Due to single nucleotide polymorphism (SNP) in different allotypes of IgG3 gene predominating in different ethnic groups, IgG3 Fc N-glycopeptide would either be identical to IgG2 Fc N-glycopeptide (in the Caucasian population)^74^ or be an isomeric coelution (undistinguishable in C18 nanoLC) of IgG4 Fc N-glycopeptide (in other ethnic groups)^59,75,76^. Given that the samples population from Cohort 2 was primarily Caucasian (11 of 14 patients), the glycopeptides were in essence resolved as IgG1, IgG2/3, and IgG4.

#### Data Analysis

The MS1 chromatogram of each sample was deconvoluted by Xtract algorithm considering 1-6 charges in FreeStyle 1.8 SP2 QF1. Deconvoluted extracted ion chromatogram (XIC) was drawn with 10 ppm error of [M+H]^+^ considering the 20 most common glycoforms and non-glycosylated peptide (NG) listed in **Supplemental Table S1**. There are multiple shared masses by different combinations of IgG subclass peptide backbones and glycoforms, which can only be distinguished by RT. The determination of RT for neutral and acid glycopeptides, and non-glycosylated peptides was done by observing the XIC profile and with the aid of Byonic v.5.7.33 search. The scan time of MS2 with Y1 and Y1F ions for each IgG subclass glycopeptide was used to help confirm the identity of peaks appeared in MS1 XIC. The determined RT ranges of each type of glycopeptides/non-glycosylated peptides are listed in **Supplemental Table S2**. Example XIC profiles with identified peaks are shown in **Supplemental Figure S2**.

### Multivariate statistical analysis

Partial Least Squares Regression (PLSR) or Discriminant Analysis (PLSDA) models were generated in MATLAB R2024a (MathWorks) using scripts developed in-house. All data input to PLSR/DA were log-transformed, centered, and scaled. The prediction accuracy of PLSDA models was determined using 5-fold cross-validation (CV), in which the data were split into separate training and testing sets using an 80:20 split. One thousand independent rounds of cross-validation were performed to determine a mean accuracy score, based on the model’s ability to predict the testing data set class labels. Statistical significance was determined by comparing this distribution of model accuracy scores against CV accuracy scores generated from 1,000 randomly permuted models with shuffled class labels using a two-sided, two-sample t-test.

A three-class PLSDA model was trained on cell type-specific Fc receptor expression frequencies to discriminate between trimester groups in Cohort 1. To compare across both cohorts, since Cohort 2 only contained samples from third trimester, PLSR models were trained on each cohort separately to predict gestational age as a continuous variable from cell type-specific Fc receptor expression frequencies. All models were orthogonalized to align the direction of maximum variation in the outcome variable (gestational age) with latent variable 1 (LV1). Univariate comparisons between model features were conducted using a non-parametric Kruskal-Wallis test followed by Dunn’s multiple comparison test.

A two-class PLSDA model was trained on the relative abundance of subclass-specific IgG glycosylation structures in matched maternal and umbilical cord samples. The least absolute shrinkage and selection operator (LASSO)^77^ was used for feature selection prior to building the model. LASSO was implemented in MATLAB as described in^78^. To ensure stability of the feature space, only features selected in >90% of LASSO iterations were used in the final PLSDA model. Pearson correlation network analysis was performed to elucidate colinear relationships with the LASSO-selected features. The resulting correlation network was trimmed to show only significant pairwise correlations (*p*<0.05).

### Unsupervised clustering of patients from clinical metadata

Dimensionality reduction and unsupervised clustering were performed in R (v.4.0). Clinical data from each cohort including race, ethnicity, fetal sex, maternal age, gravidity, pregravid body mass index (BMI) and Edinburgh Postnatal Depression Scale (EPDS) scores were extracted from patient charts. CRP was measured in maternal plasma sampled at the time of delivery using an ELISA kit (Invitrogen KHA0031, 1:30,000 dilution). Patients with incomplete observations were removed from the matrix. Data from Cohort 1 were log-transformed, centered, and scaled prior to dimensionality reduction. PCA was used for dimensionality reduction and k-means clustering was performed on PC1 and PC2 scores. K=3 clusters was selected based on the within-sample-similarity (WSS) criterion. To validate these clusters, data from Cohort 2 were projected into the principal component space by multiplying the log-transformed, centered, and scaled features from Cohort 2 by the feature loadings matrix from Cohort 1. Cohort 2 patients were assigned to one cluster by identifying the minimum Euclidean distance between the sample projection in latent space and the cluster centroids.

### Statistical analysis

Statistical analysis was performed in R (v.4.0) and MATLAB R2024a. A Mann-Whitney rank sum test was used for two-group comparisons. Paired comparisons (i.e., comparing expression of two Fc receptors within the same patient) were performed with the Wilcoxon signed rank test. The Kruskal-Wallis test (non-parametric) with a Dunn’s test for multiple comparisons or an ANOVA test (parametric) with Tukey’s test for multiple comparisons was performed for analyses involving three or more groups. Partial correlation network analyses were performed by controlling for gestational age and maternal covariates (age, gravidity, pregravid BMI, and EPDS). Correlations with R > 0.15 and *p*-value < 0.1 were shown in the resulting networks.

## Acknowledgements

This work was funded by the NIH NIAID (1R01AI184565-01), the Jeffress Trust Award Program in Research Advancing Health, and the Hypothesis Fund awarded to SD, and NIH R24GM137782 and NSF GlycoMIP DMR-1933525 awarded to PA. REW received support from the NIH NIGMS (T32-GM145443), the Philanthropic Educational Organization (PEO) Scholar Award Fund, and the University of Virginia School of Engineering and Applied Sciences Endowed Graduate Fellowship. We thank the Molecular, Immunologic, and Translational Sciences (MITS) Core Facility at UVA (CCSG grant number 2P30CA044579-26) for assistance with multiplex IHC experiments and data analysis, the Biospecimen Tissue Repository and Foundation (BTRF) at UVA for providing access to clinical specimens, the Flow Cytometry Core Facility (FCCF) at UVA for performing Luminex experiments, and the Center for Complex Carbohydrate Research (CCRC) at the University of Georgia for assistance with glycoproteomic analysis (R24GM137782, PI: Parastoo Azadi). We thank Drs. Robert Fuller and Christopher Chisholm and Ms. Amanda Urban for support with the clinical study at UVA and members of the Dolatshahi Lab for helpful feedback. Finally, we thank the patient donors who participated in this study.

## Author Contributions

REW, DJD, and SD were responsible for the conceptualization of the project and methodologies. AB, GM, VK, LZ, DJD, and YA managed the clinical cohorts. REW, AB, AS, CS, ISM, PA, and XY conducted experiments. ISM and PA provided guidance and supervision on multiplex immunofluorescence and glycoprotoeomics, respectively. REW performed data analysis. REW and SD wrote the manuscript with input from all authors. REW, AB, GM, VK, AS, CS, XY, PA, ISM, LZ, DJD, YA, and SD read and approved the final version of the manuscript.

## Data Availability Statement

Processed data from mIHC, glycoproteomics, and immunoglobulin isotyping experiments along with associated metadata have been made publicly available on University of Virginia Libra Dataverse (https://doi.org/10.18130/V3/UKEWUQ). Codes for multivariate data analysis are available on GitHub (https://github.com/Dolatshahi-Lab/PLSR-DA). Raw images and data will be made available upon reasonable request.

## Declaration of Interests

The authors declare no competing interests. Yalda Afshar is a consultant to BilliontoOne and J&JMedTech, unrelated to the work presented here.

---

**Supplemental Figure S1.**
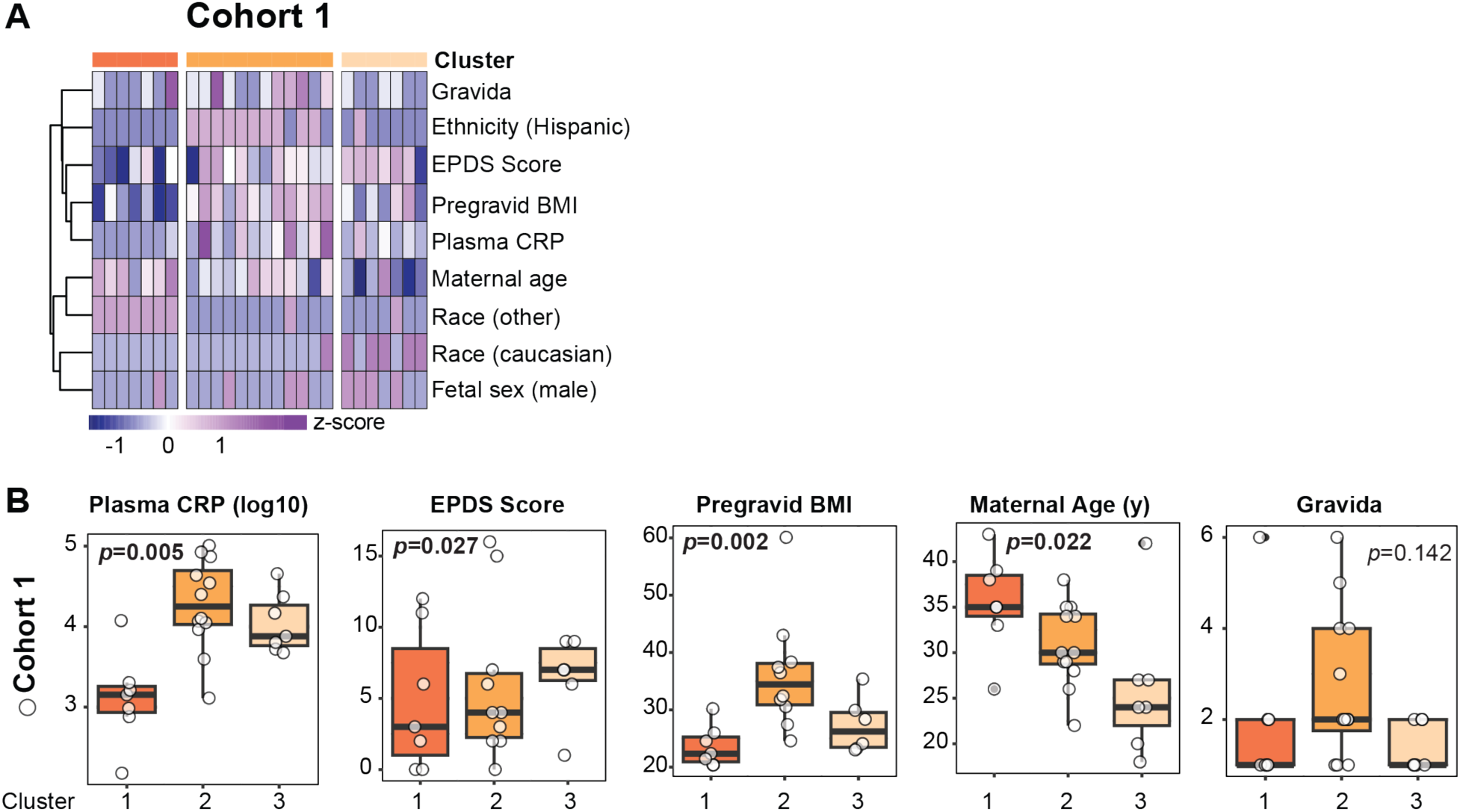
Clinical covariates across patient clusters. (A) Heatmaps show z-scored values of each feature used for PCA of Cohort 1 (n=26). Rows are clustered by hierarchical clustering. Columns are organized by cluster identity, corresponding to the PCA scores plots in Figure 1. (B) Continuous-valued clinical features that were included in the PCA model. Significance was determined by a Kruskal-Wallis test (*p*<0.1 are emphasized in boldface font).

**Supplemental Figure S2.**
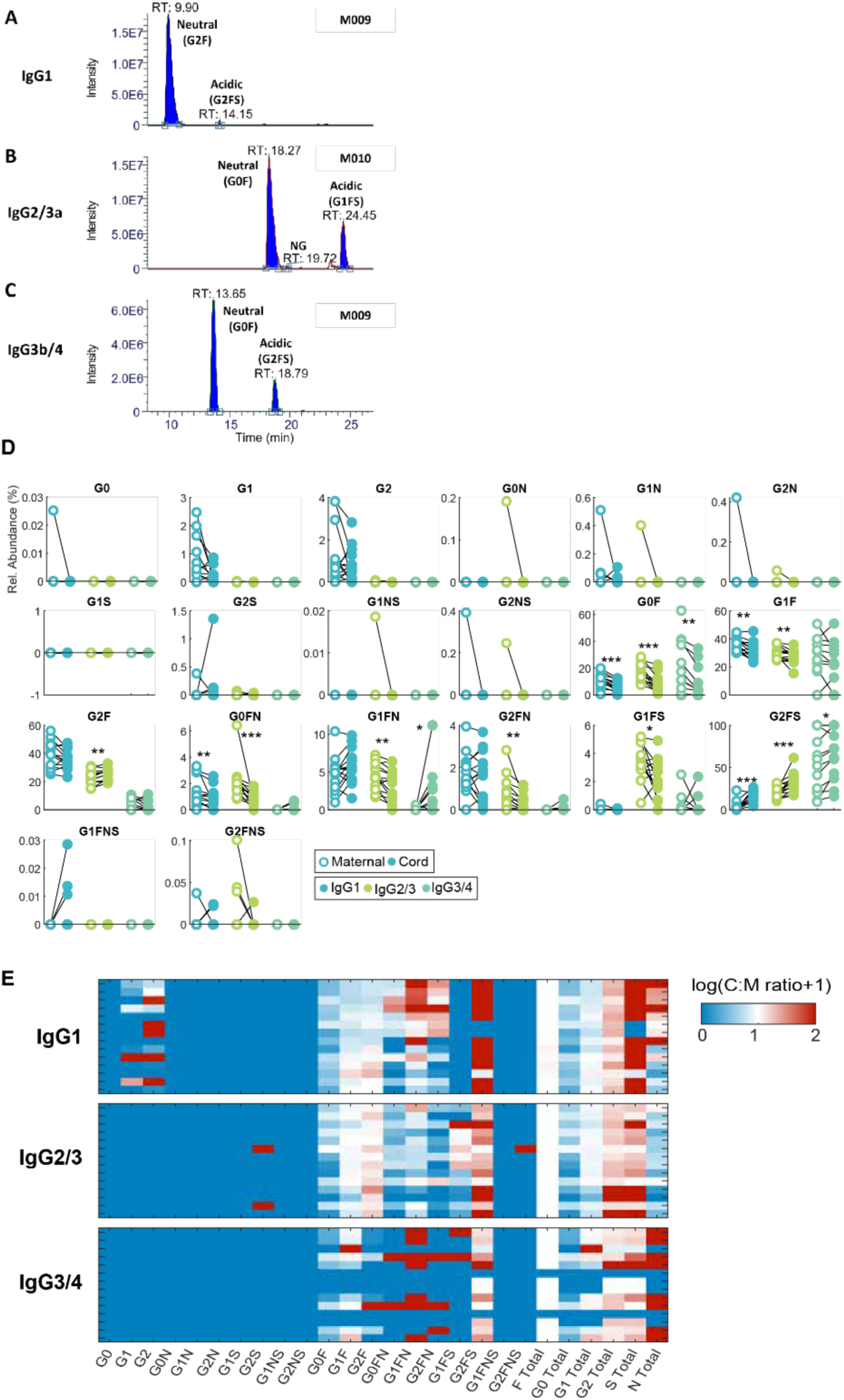
Subclass-specific Fc glycan analysis by liquid chromatography mass spectrometry (LC-MS). (A-C) Example XIC profiles for glycopeptides/non-glycosylated peptides of IgG1 (A), IgG2/3 (B), and IgG3/4 (C). Panels A and C were from sample M009, B was from sample M010. (D) Paired comparisons between maternal and cord relative abundance of subclass-specific Fc glycans. (\**p*<0.05, \*\**p*<0.01, \*\*\**p*<0.001, Wilcoxon signed rank test). (E) Heatmap shows the log-transformed transfer ratio of subclass-specific IgG glycans. Each row represents one maternal-cord dyad.

**Supplemental Figure S3.**
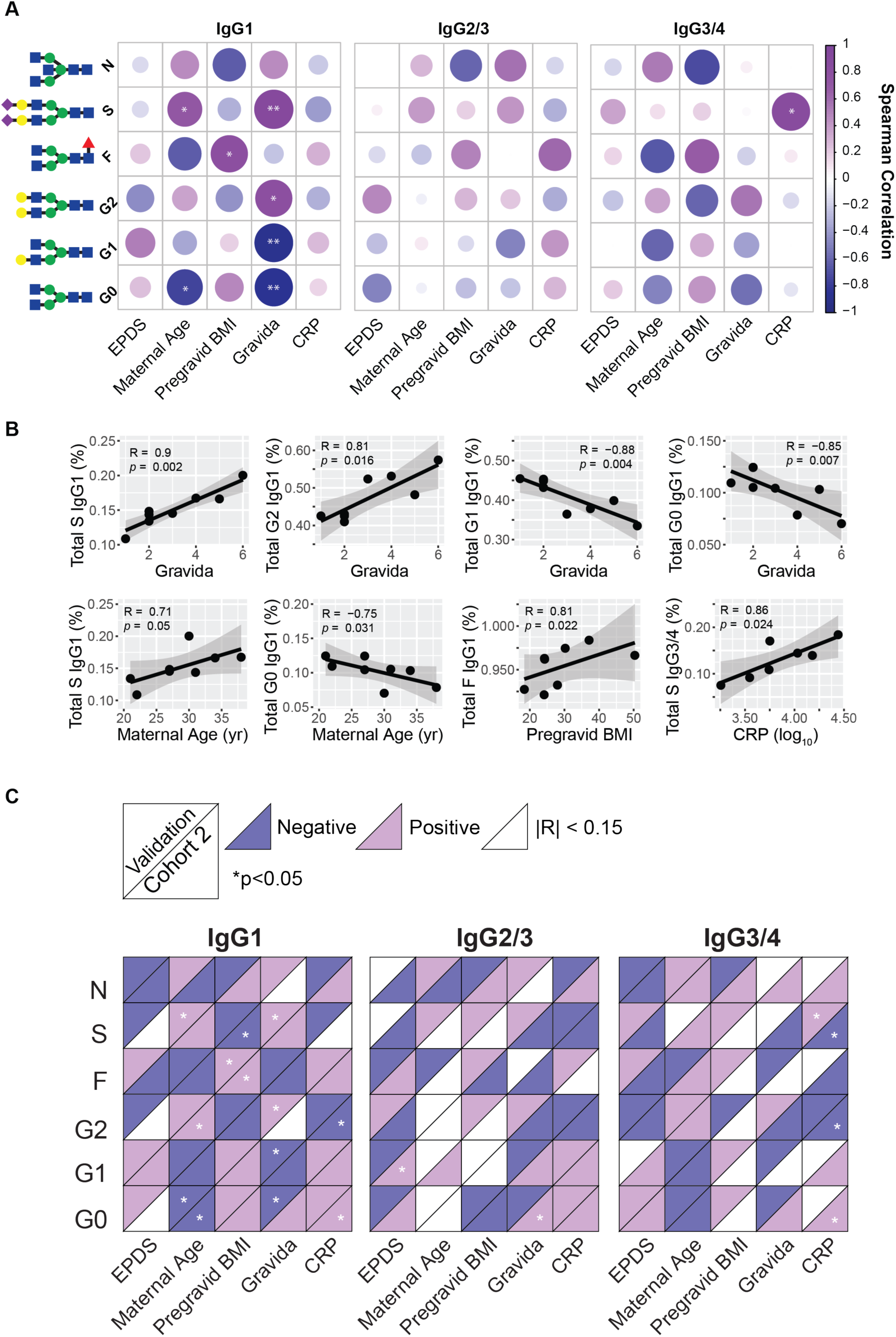
Validation of correlations between clinical covariates and maternal subclass-specific IgG glycans in a separate cohort. (A) Correlation heatmap between clinical metadata and summary glycans in a second patient sample. (\**p*<0.05, \*\**p*<0.01, Spearman correlation). (B) Scatter plots show significant correlations from (A). Spearman correlation coefficient and corresponding *p*-value are labeled on the axes. (C) Qualitative comparison of results from two sample populations. For each box, the top triangle displays the directionality of correlations in the validation cohort, and the bottom triangle displays directionality of correlations in Cohort 2 (corresponding to the data in Figure 2). Correlations with |R| < 0.15 are colored white. Significant correlations are marked (\**p*<0.05).

**Supplemental Figure S4.**
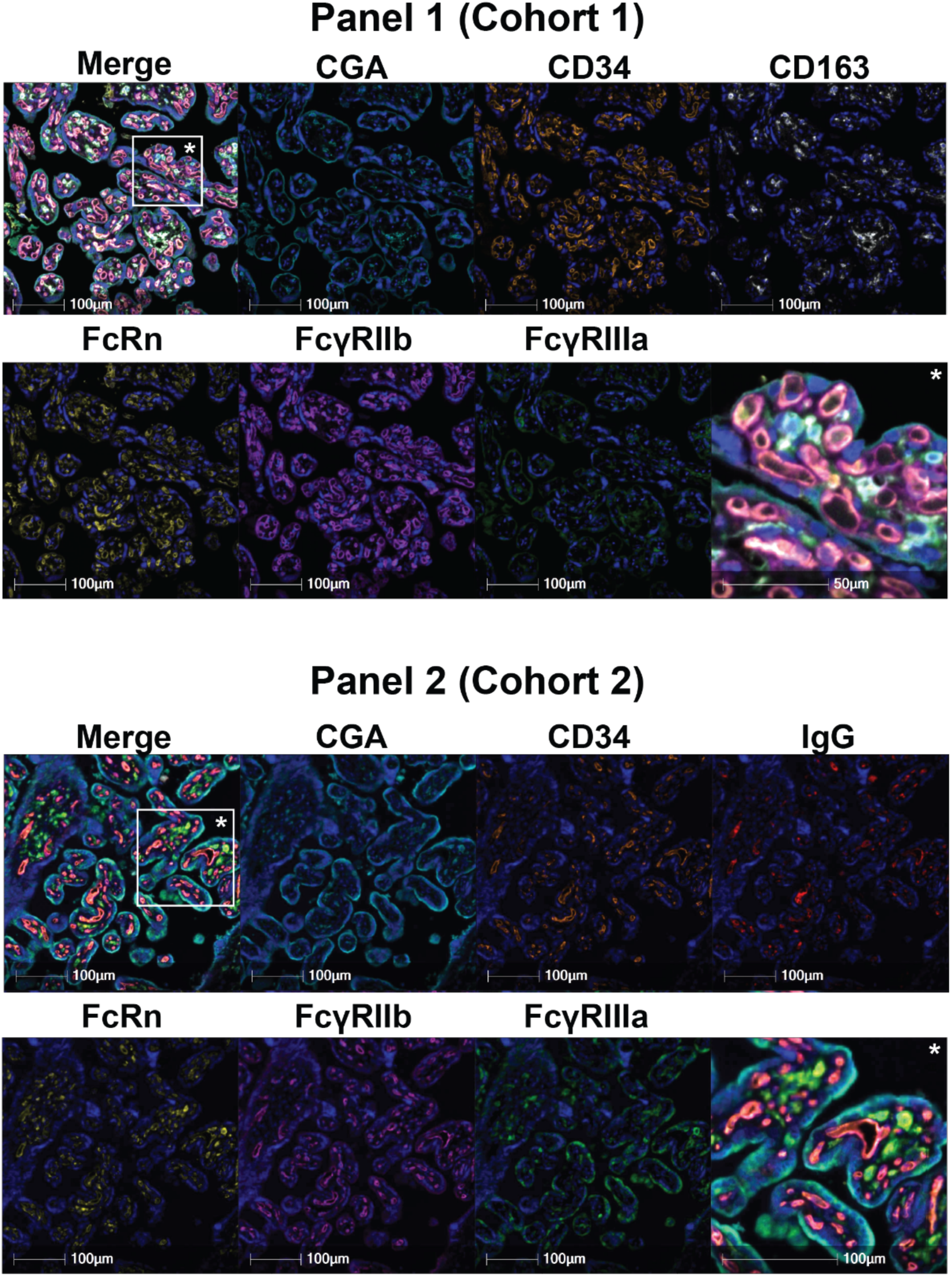
Representative mIHC images from panel 1 and panel 2. Representative single- and multiplex images from panel 1 (top) and panel 2 (bottom). Each single-plex image is labeled. The asterisk (*) indicates a portion of the image shown at higher magnification in the bottom right square of each image. Scale bars, 100 μm.

**Supplemental Figure S5.**
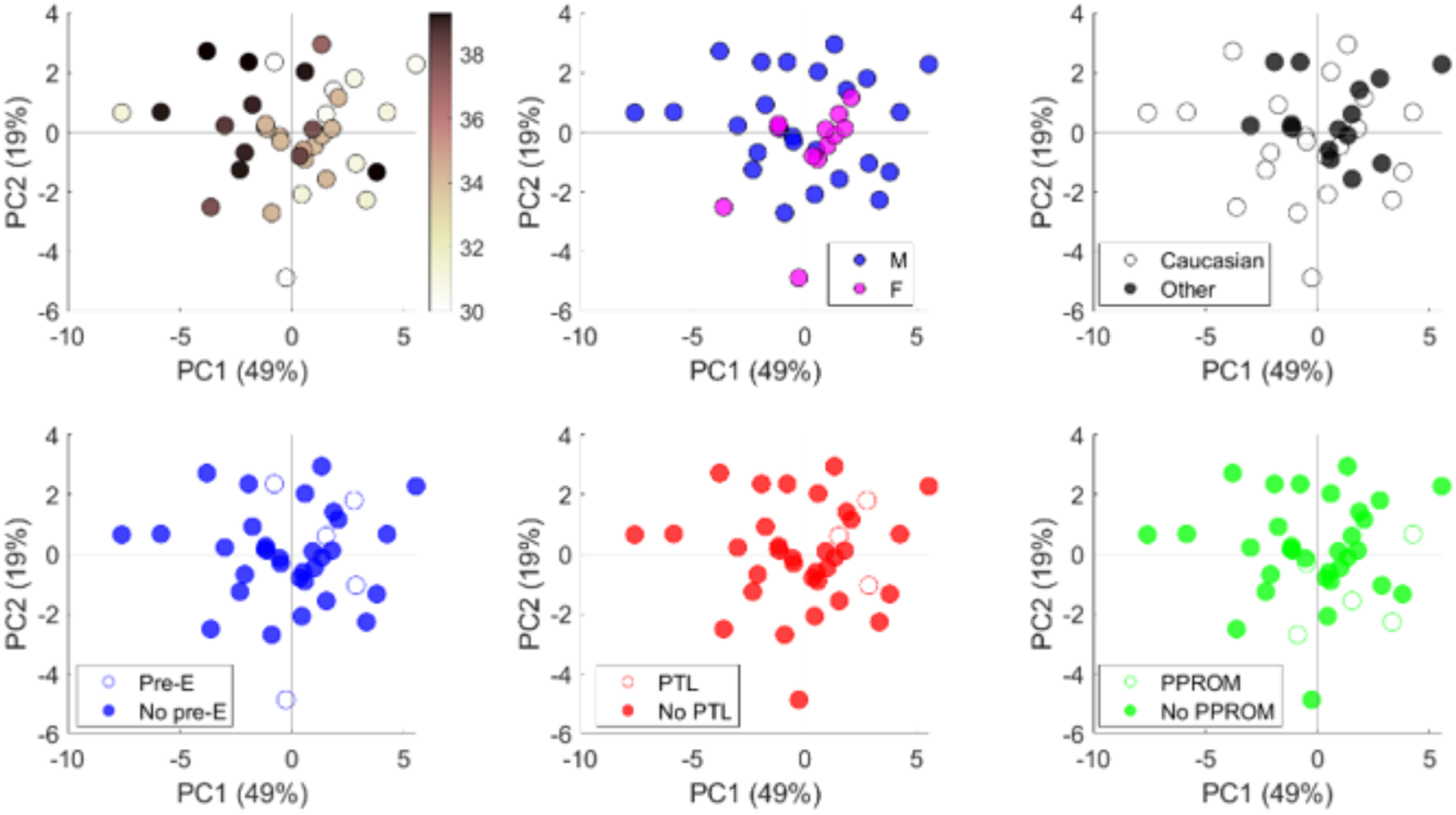
The indication of preterm delivery does not affect placental Fc receptor expression. PCA scores plots of patients who delivered prematurely (<37 weeks) in Cohort 2 colored by gestational age, fetal sex, maternal race, or indications of preterm birth (preeclampsia, Pre-E, preterm labor, PTL, or premature preterm rupture of membranes, PRROM).

**Supplemental Figure S6.**
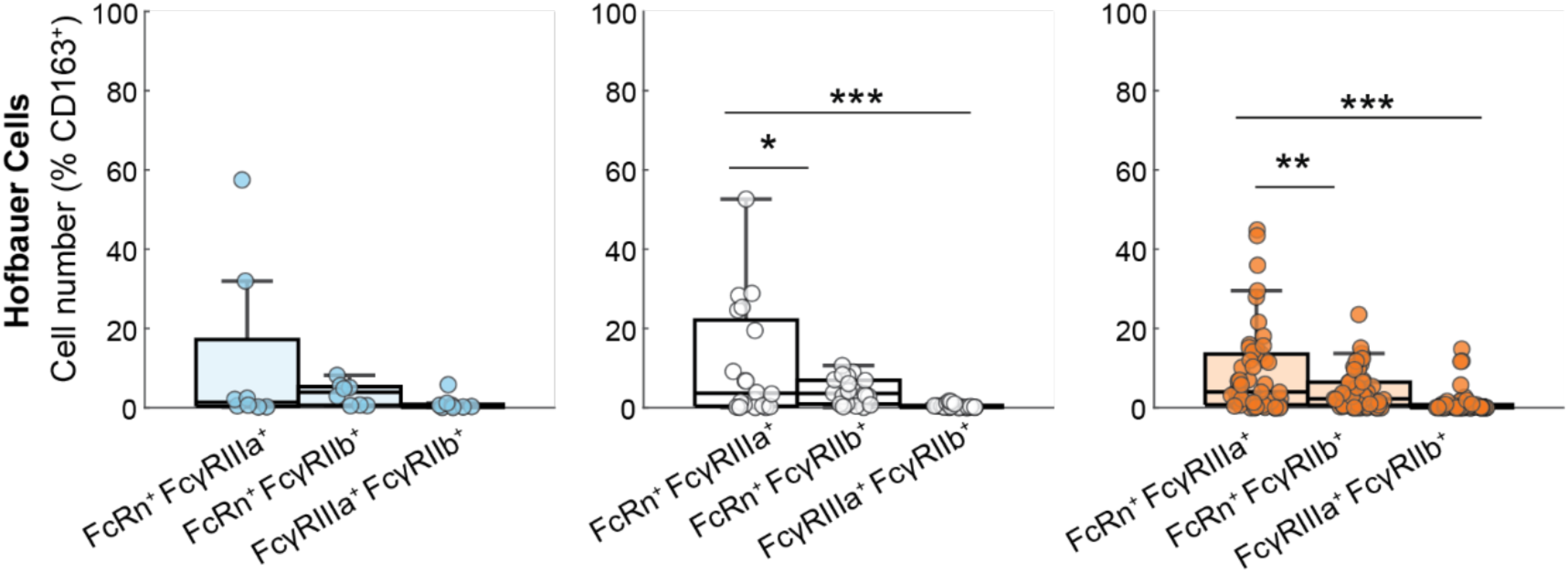
Pairwise colocalization analysis of Fc receptors expressed by Hofbauer cells. Boxplots show pairwise colocalizations of Fc receptors on Hofbauer cells, expressed as the average percentage of total Hofbauer cells in each image from each patient. Patients from trimesters are shown on separate axes (blue, 1^st^ trimester; white, 2^nd^ trimester; orange, 3^rd^ trimester). (\*\*\**p*<0.001, \*\**p*<0.01, \**p*<0.05, ANOVA with Tukey’s test for multiple comparisons).

**Supplemental Figure S7.**
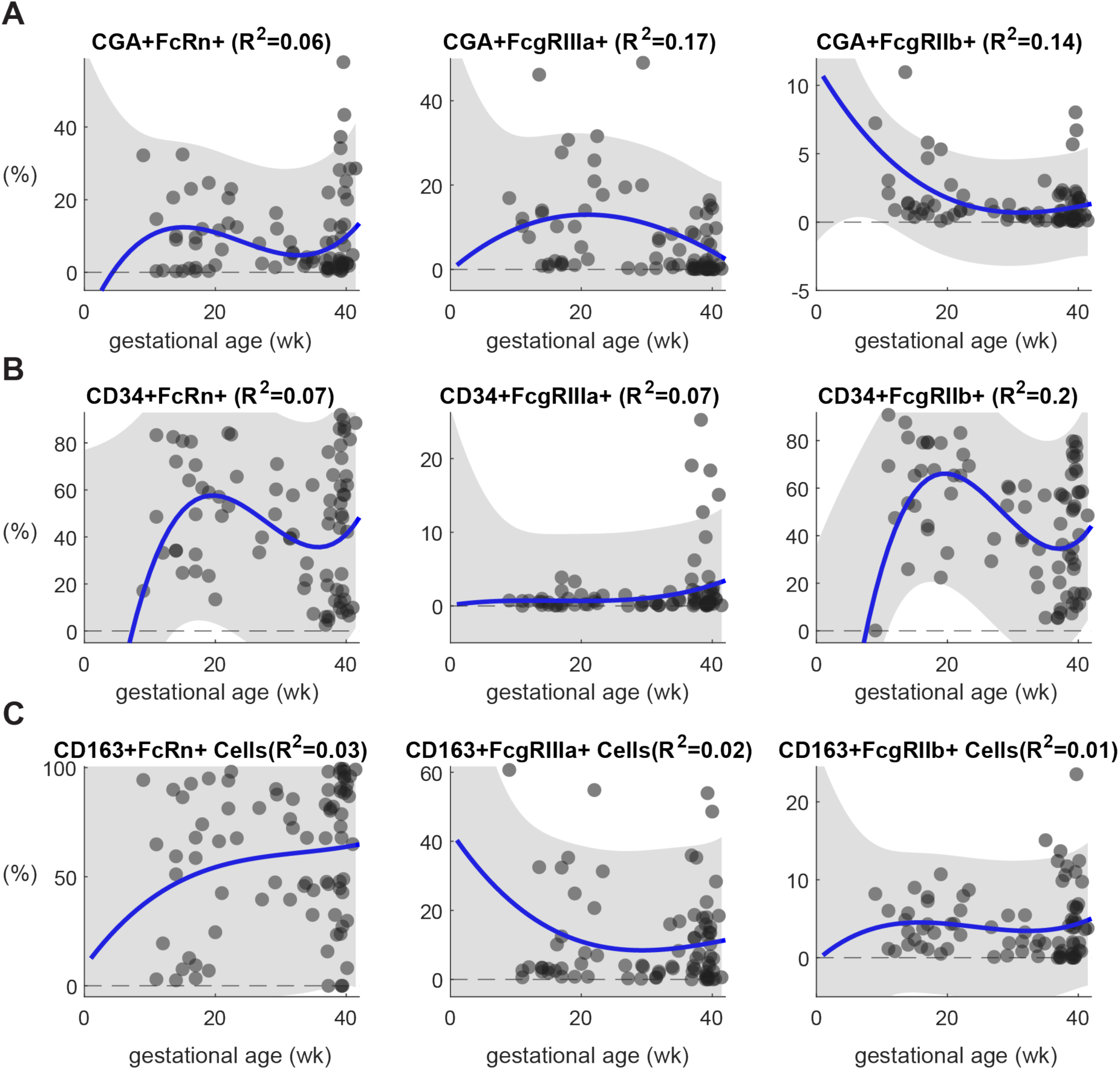
Polynomial regression analysis of cell type-specific Fc receptor expression across gestation in Cohort 1. (A-C) Scatter plots showing inferred gestational expression trajectories for each receptor expressed on STBs (A), ECs (B), and HBCs (C) in Cohort 1. Each point represents the average expression frequency from a single patient tissue. The solid lines represent a 3^rd^-order polynomial curve fit. The gray shaded region denotes 95% confidence intervals. The regression R^2^ is listed in the top right of each coordinate plane.

**Supplemental Figure S8.**
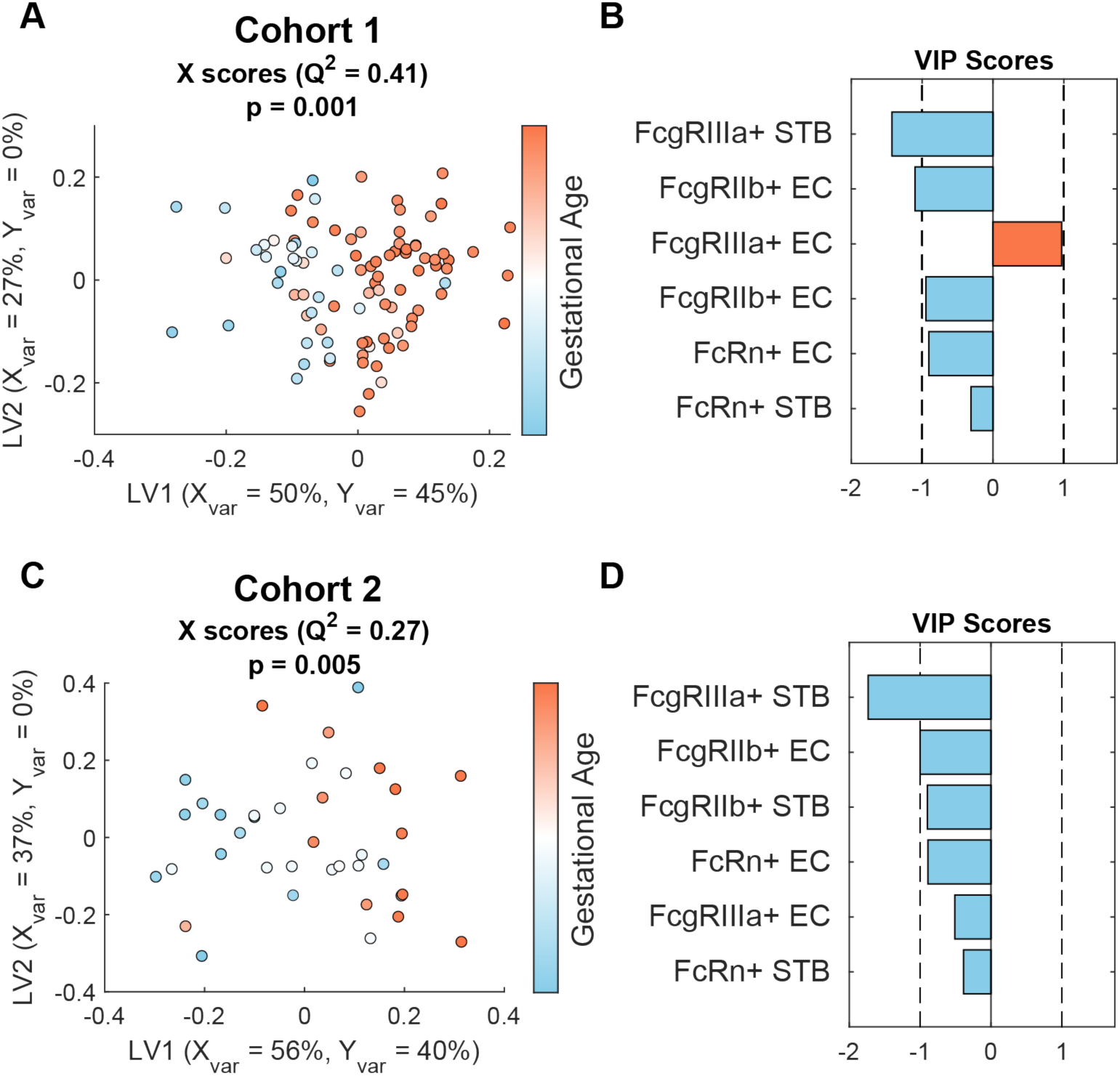
Similar temporal trends in cell type specific Fc receptor expression observed across cohorts. OPLSR models trained on Fc receptor expression data from Cohort 1 (A-B) and cohort (C-D). (A,C) X scores plot, where points are colored according to gestational age. Each point represents one patient tissue. Q^2^ is a metric of the model’s prediction performance, and the p-value was determined by permutation testing by comparing Q^2^ from this model against 1,000 models with randomly shuffled Y-labels. (B,D) VIP scores bar plot. The model trained on data from Cohort 1 (A-B) was used to predict gestational age in Cohort 2, as shown in Figure 5.

**Supplemental Figure S9.**
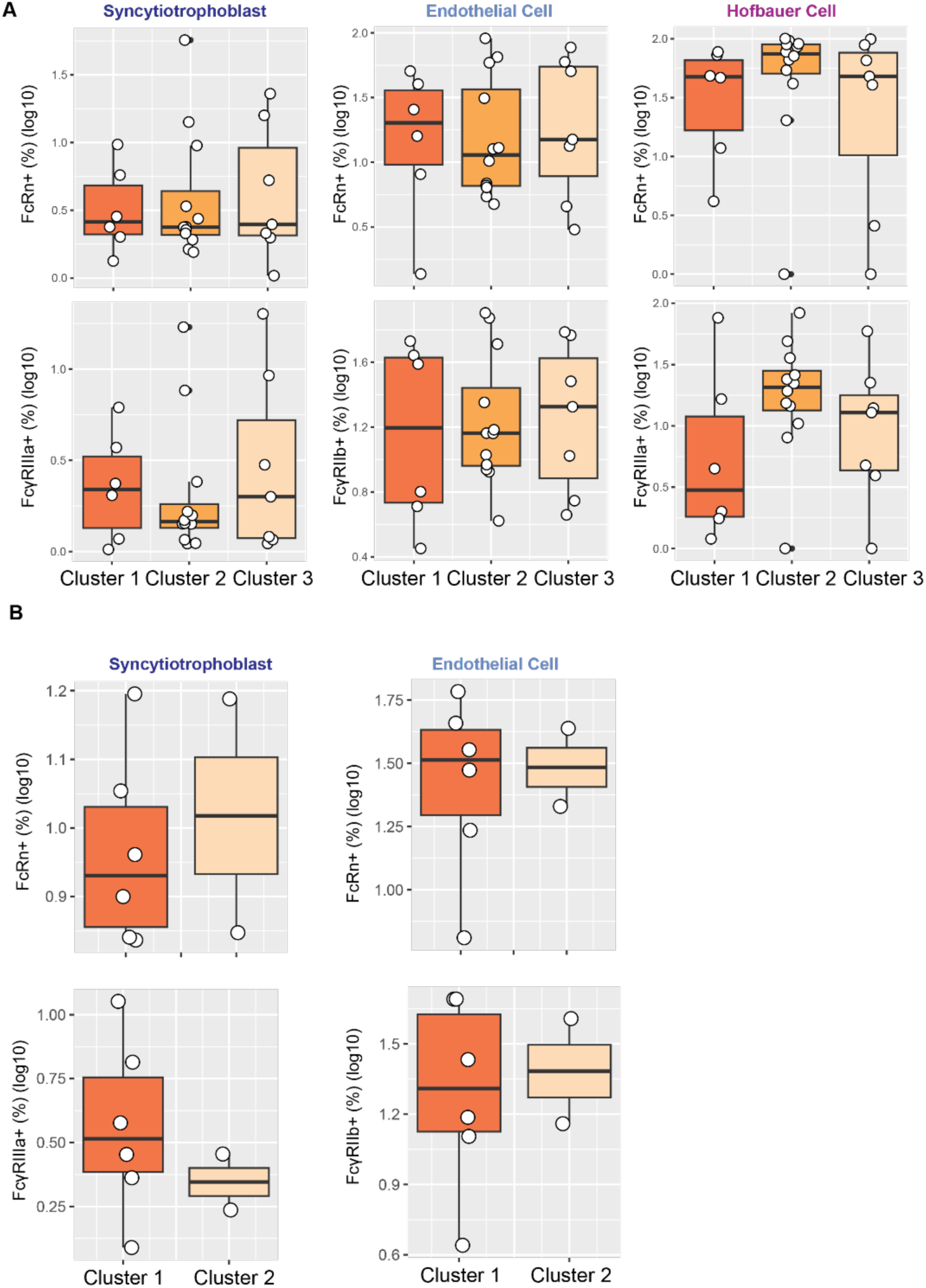
Fc receptor expression across patient clusters. (A) Cell type-specific Fc receptor expression levels across clusters in Cohort 1, related to Figure 1. No comparisons were found to be significantly different at (p<0.1) (Kruskal-Wallis test with Dunn’s test for multiple comparisons). (B) Cell type-specific Fc receptor expression levels across clusters in Cohort 2, related to Figure 1. No comparisons were found to be significantly different at (p<0.1) (Kruskal-Wallis test with Dunn’s test for multiple comparisons). Only clusters 1 and 3 are shown because no patients in cluster 2 had matched placental tissue and plasma available.

**Supplemental Figure S10.**
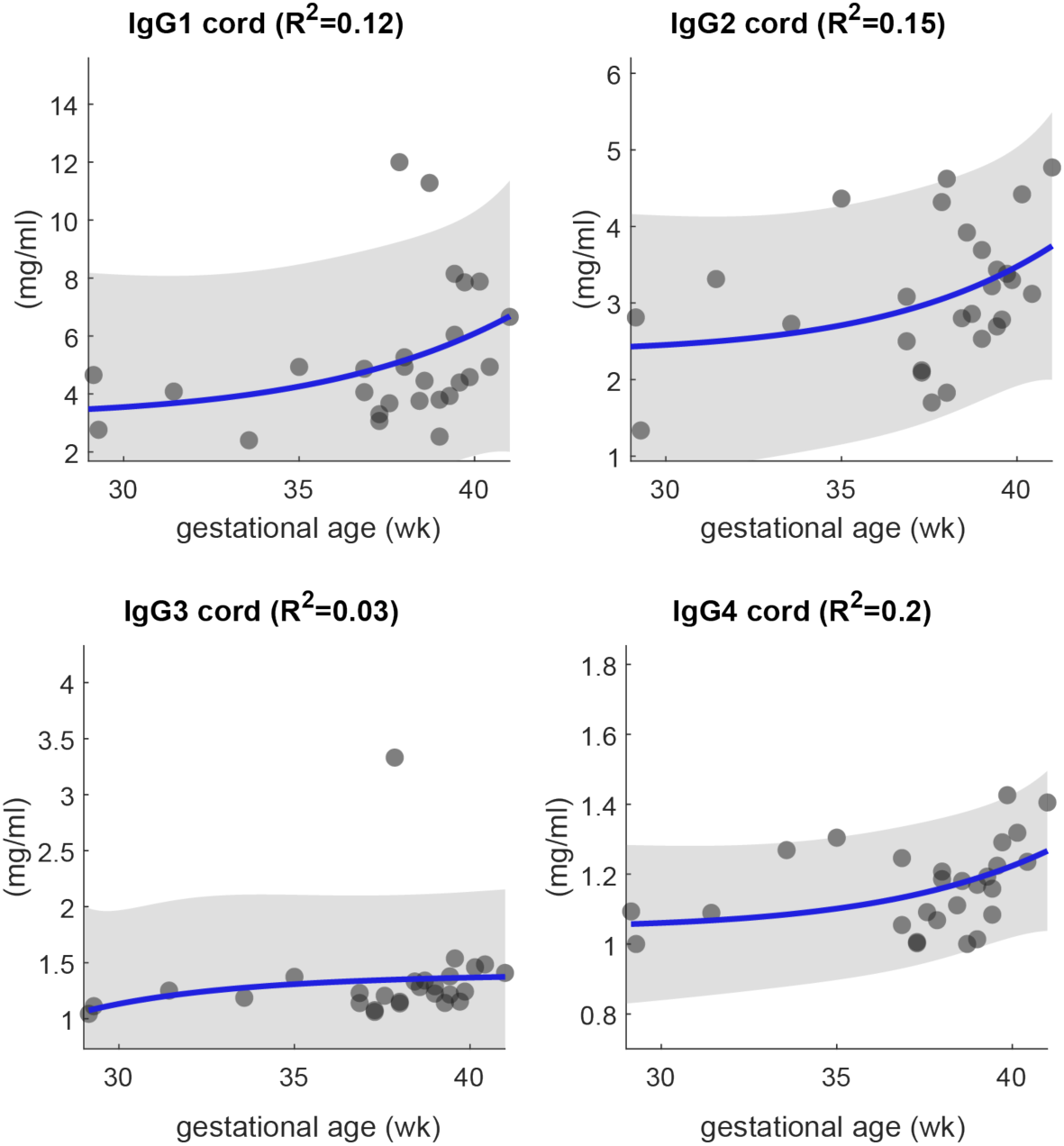
IgG transfer is dynamically regulated across third trimester. Scatter plots show the concentration of umbilical cord IgG subclasses as a function of gestational age in Cohort 1. The solid lines are exponential regressions with 95% confidence intervals shaded in gray. Regression R^2^ are shown in each figure title.

**Supplemental Figure S11.**
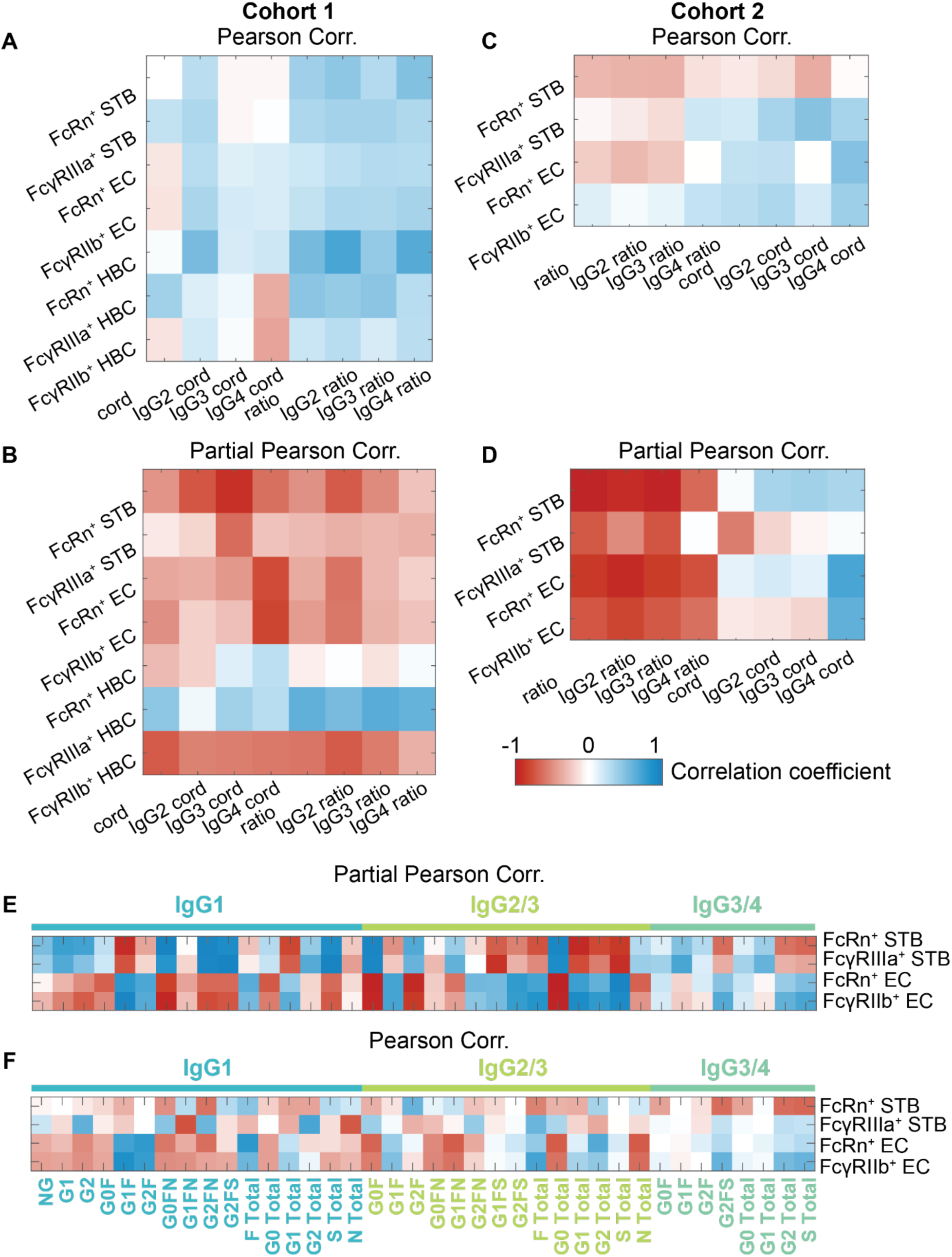
Controlling for clinical covariates reveals positive associations between Fc receptor expression frequencies and IgG subclass transfer efficiency. (A-D) Heatmaps show Pearson correlation coefficients (A,C) and partial Pearson correlation coefficients (B,D) between cell type-specific Fc receptor expression frequencies and IgG subclass cord concentrations or cord:maternal ratios after controlling for clinical covariates used for clustering analysis in Figure 1. Data from Cohort 1 are shown in (A,B) and data from Cohort 2 are shown in (C,D). (E,F) Heatmaps show Pearson correlation coefficients (E) and partial Pearson correlation coefficients (F) between cell type-specific Fc receptor expression frequencies and subclass-specific glycoforms after controlling for clinical covariates used for clustering analysis in Figure 1. The colored bar denotes the subclass-specific glycoforms. Glycoforms were omitted from the heatmap if they fell below the limit of detection, resulting in NA correlation values with all Fc receptors.

**Supplemental Table 1.**
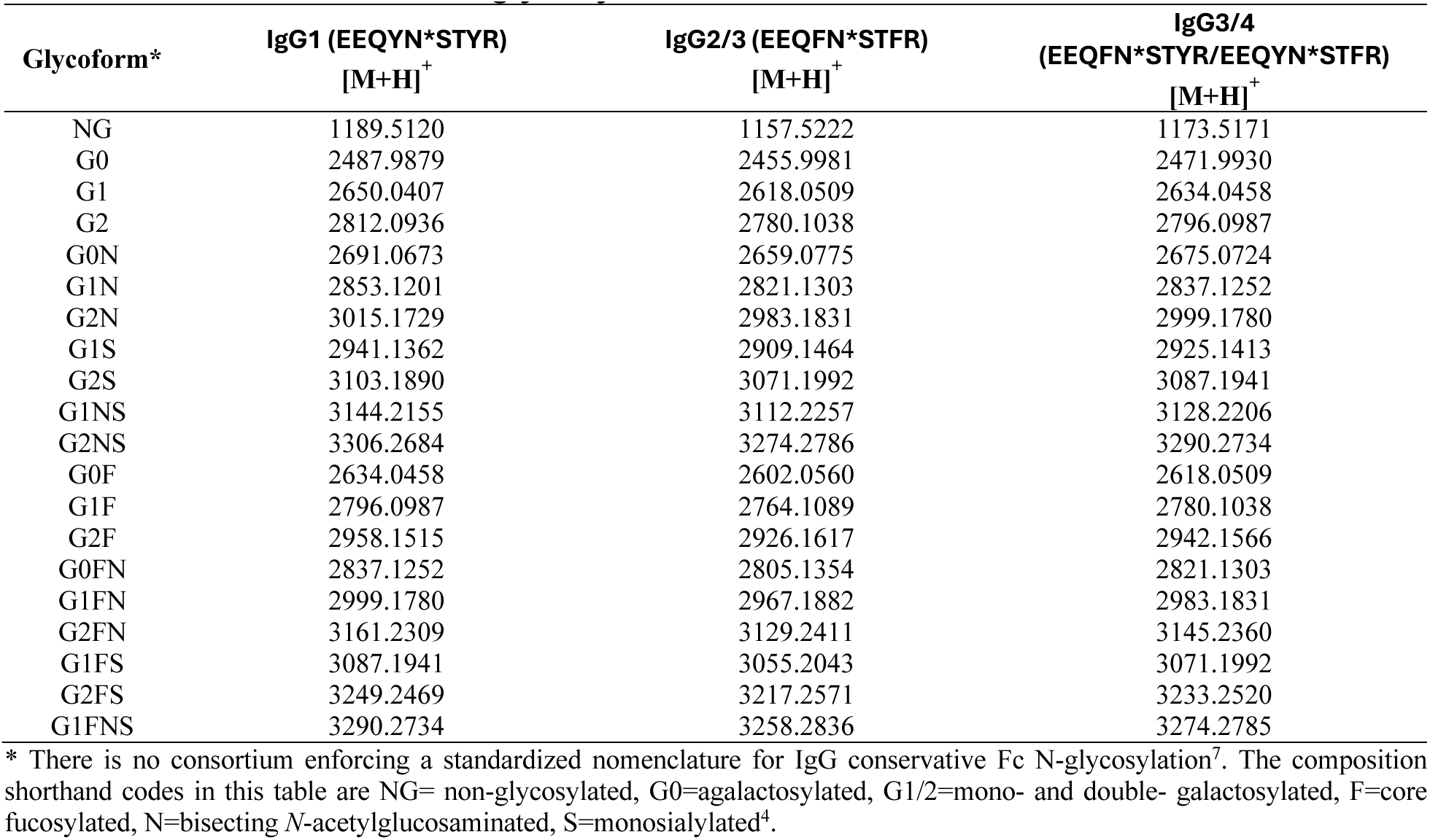
Intact glycopeptide/non-glycosylated peptide masses of IgG subclasses Fc conservative N-glycosylation.

**Supplemental Table 2.**
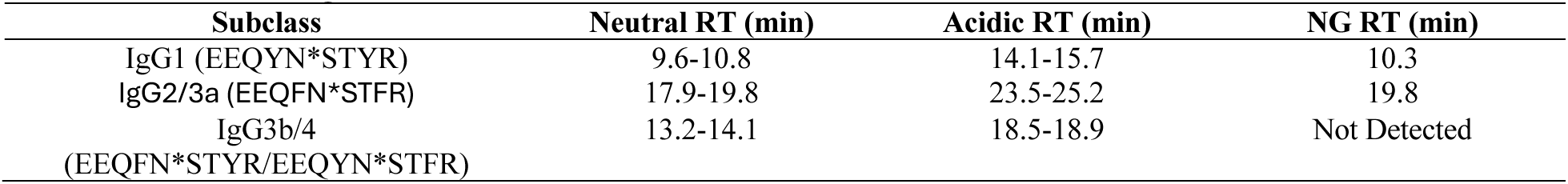
Retention time ranges for glycopeptides/non-glycosylated peptides from all IgG subclasses.

## Notes

https://dataverse.lib.virginia.edu/dataset.xhtml?persistentId=doi:10.18130/V3/UKEWUQ

https://github.com/Dolatshahi-Lab/PLSR-DA

