## Supplemental Materials (Figures, Tables) for "Multi-factorial regulatory networks of placental antibody transfer by Fc receptors and maternal IgG Fc characteristics are modulated by clinical covariate profiles"

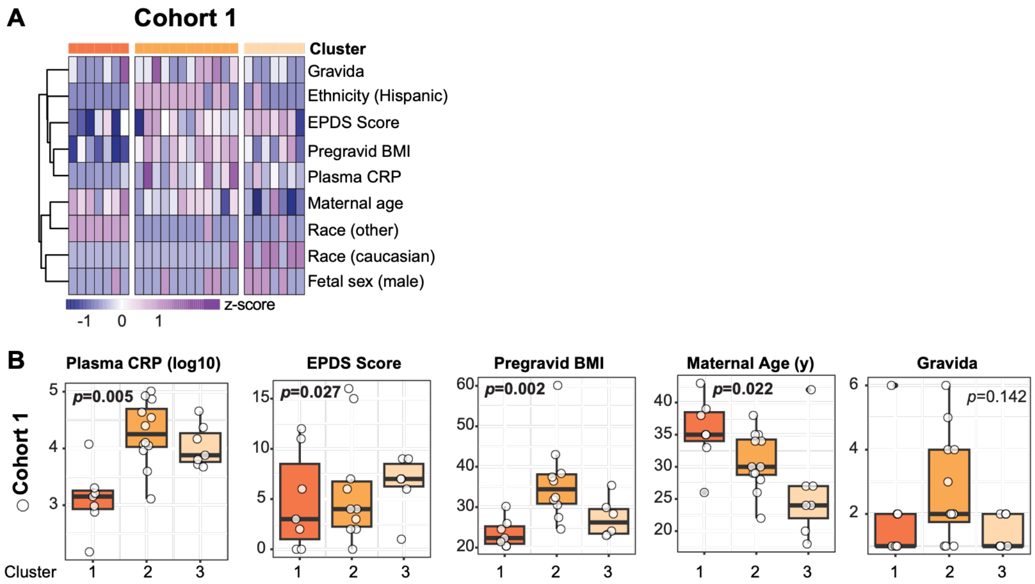


***Supplemental Figure S1. Clinical covariates across patient clusters.***

(A) Heatmap shows z-scored values of each feature used for PCA of Cohort 1. Rows are clustered by hierarchical clustering. Columns are organized by cluster identity, corresponding to the PCA scores plot in Figure 1E. (B) Continuous-valued clinical features from the PCA model compared across clusters. Significance was determined by a Kruskal-Wallis test (*p*<0.05 are emphasized in boldface font).

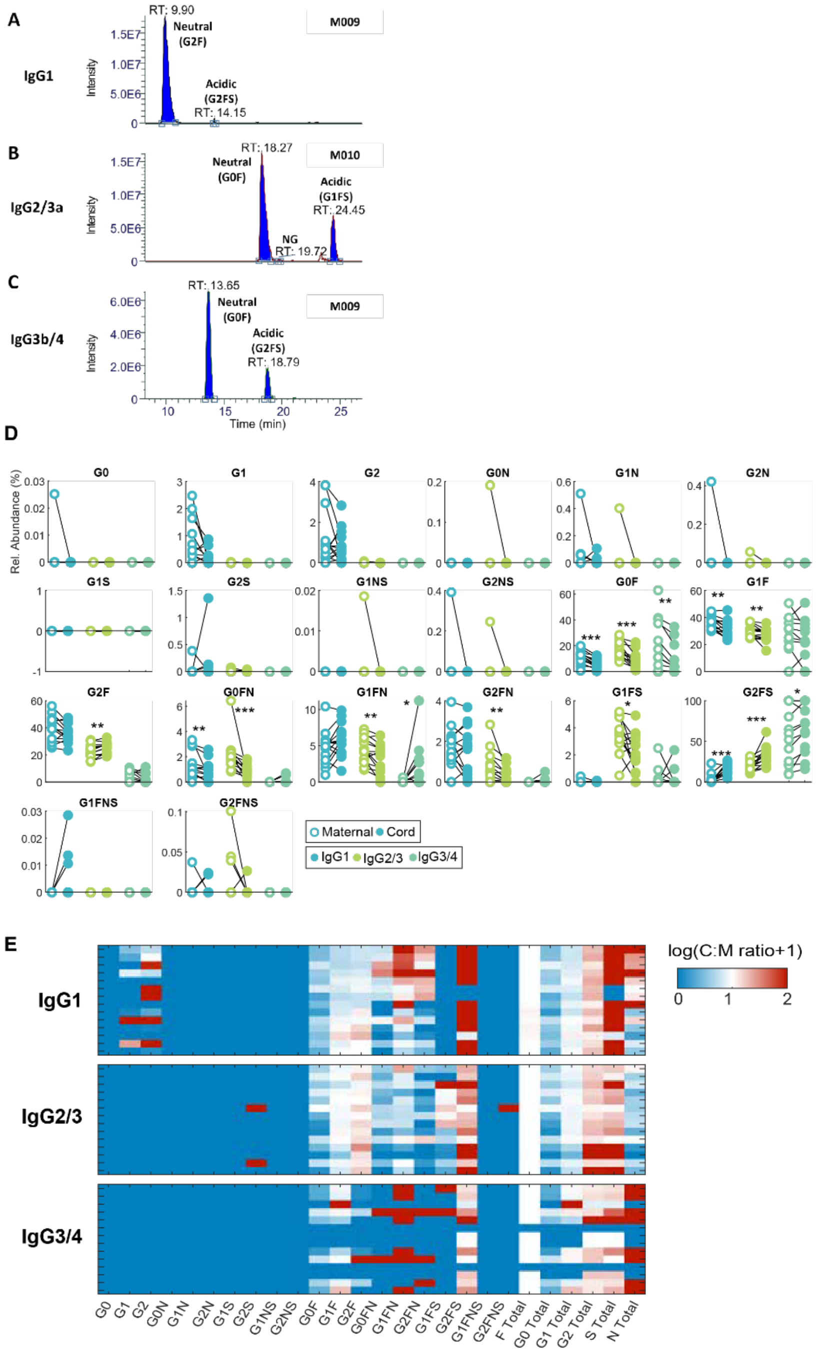


***Supplemental Figure S2. Subclass-specific Fc glycan analysis by liquid chromatography mass spectrometry (LC-MS).***

(A-C) Example XIC profiles for glycopeptides/non-glycosylated peptides of IgG1 (A), IgG2/3 (B), and IgG3/4 (C). Panels A and C were from sample M009, B was from sample M010. (D) Paired comparisons between maternal and cord relative abundance of subclass-specific Fc glycans. (**p*<0.05, ***p*<0.01, ****p*<0.001, Wilcoxon signed rank test). (E) Heatmap shows the log-transformed transfer ratio of subclass-specific IgG glycans. Each row represents one maternal-cord dyad.

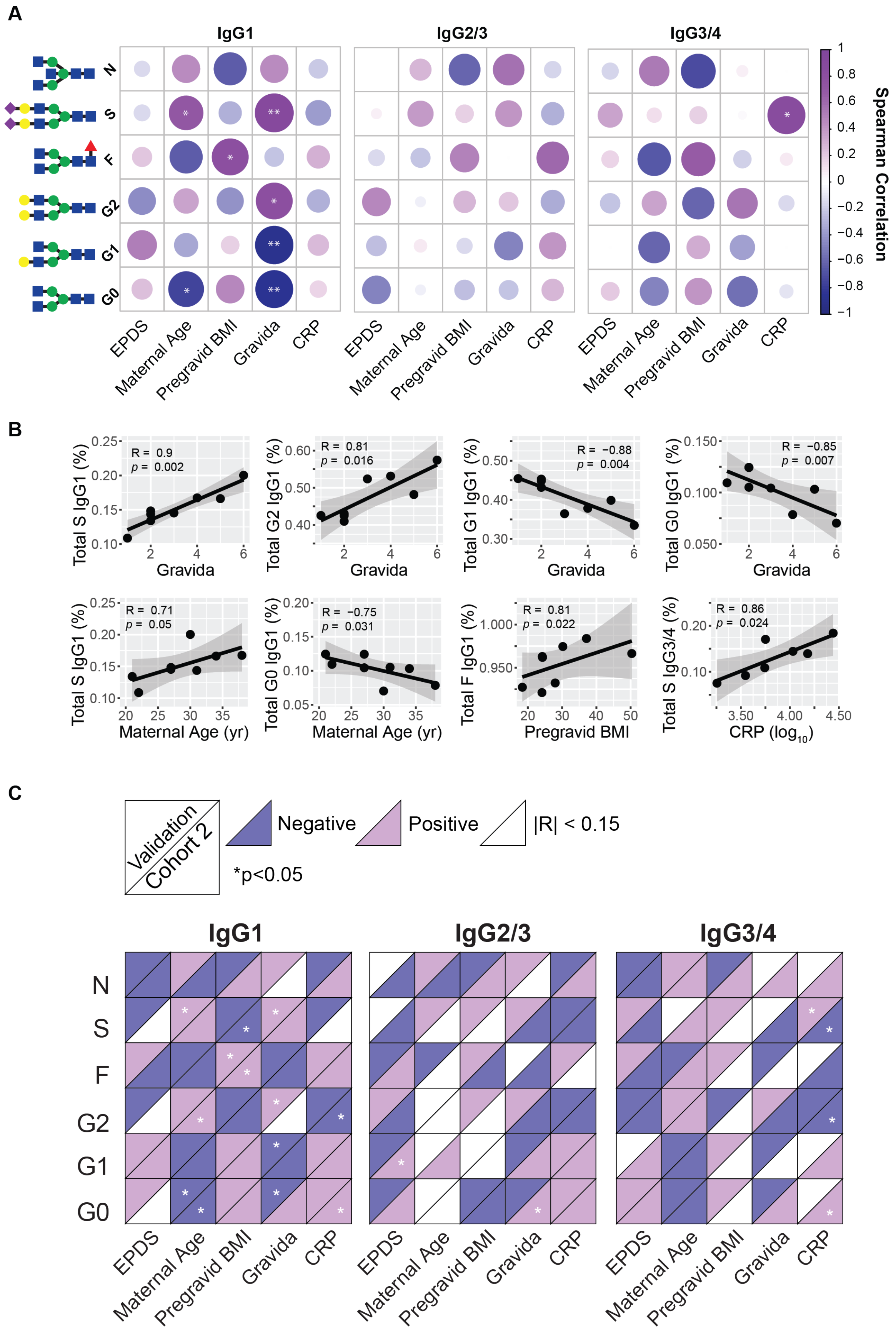
A
IgG1
IgG2/3
IgG3/4
N
S
F
G2
G1
G0
EPDS
Maternal Age
Pregravid BMI
Gravida
CRP
Spearman Correlation
B
R = 0.9
p = 0.002
Total S IgG1 (%)
Gravida
R = 0.81
p = 0.016
Total G2 IgG1 (%)
R = −0.88
p = 0.004
Total G1 IgG1 (%)
R = −0.85
p = 0.007
Total G0 IgG1 (%)
R = 0.71
p = 0.05
Maternal Age (yr)
R = −0.75
p = 0.031
R = 0.81
p = 0.022
Total F IgG1 (%)
Pregravid BMI
R = 0.86
p = 0.024
Total S IgG3/4 (%)
CRP (log10)
C
Validation
Cohort 2
Negative
Positive
|R| < 0.15
*p<0.05

***Supplemental Figure S3. Validation of correlations between clinical covariates and maternal subclass-specific IgG glycans in a separate cohort.***

(A) Correlation heatmap between clinical metadata and summary glycans in a second patient sample. (*$p<0.05$, **$p<0.01$, Spearman correlation). (B) Scatter plots show significant correlations from (A). Spearman correlation coefficient and corresponding *p*-value are labeled on the axes. (C) Qualitative comparison of results from two sample populations. For each box, the top triangle displays the directionality of correlations in the validation cohort, and the bottom triangle displays directionality of correlations in Cohort 2 (corresponding to the data in Figure 2). Correlations with $|R| < 0.15$ are colored white. Significant correlations are marked (*$p<0.05$).

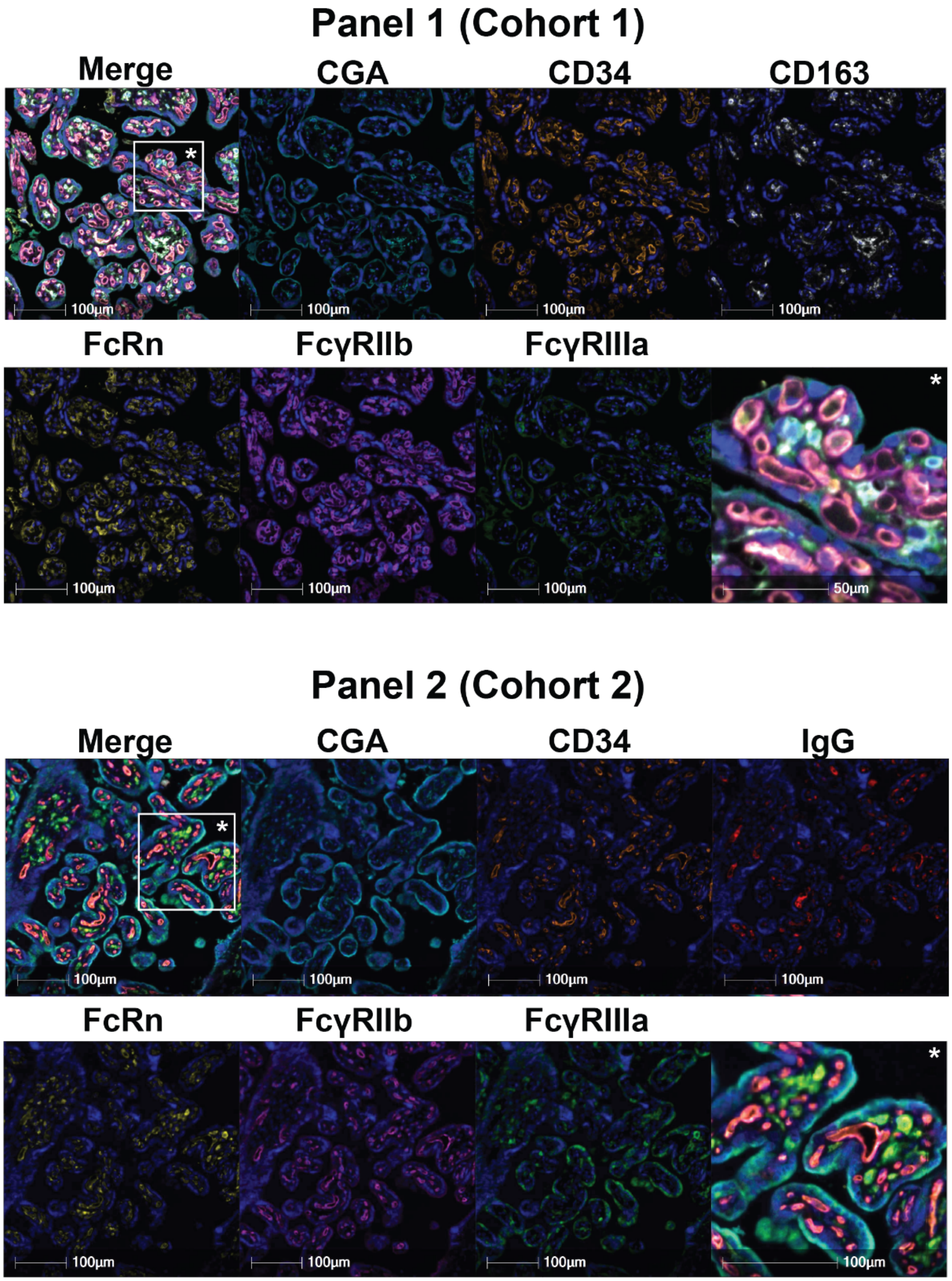


***Supplemental Figure S4. Representative mIHC images from panel 1 and panel 2.***

Representative single- and multiplex images from panel 1 (top) and panel 2 (bottom). Each single-plex image is labeled. The asterisk (*) indicates a portion of the image shown at higher magnification in the bottom right square of each image. Scale bars, 100 µm.

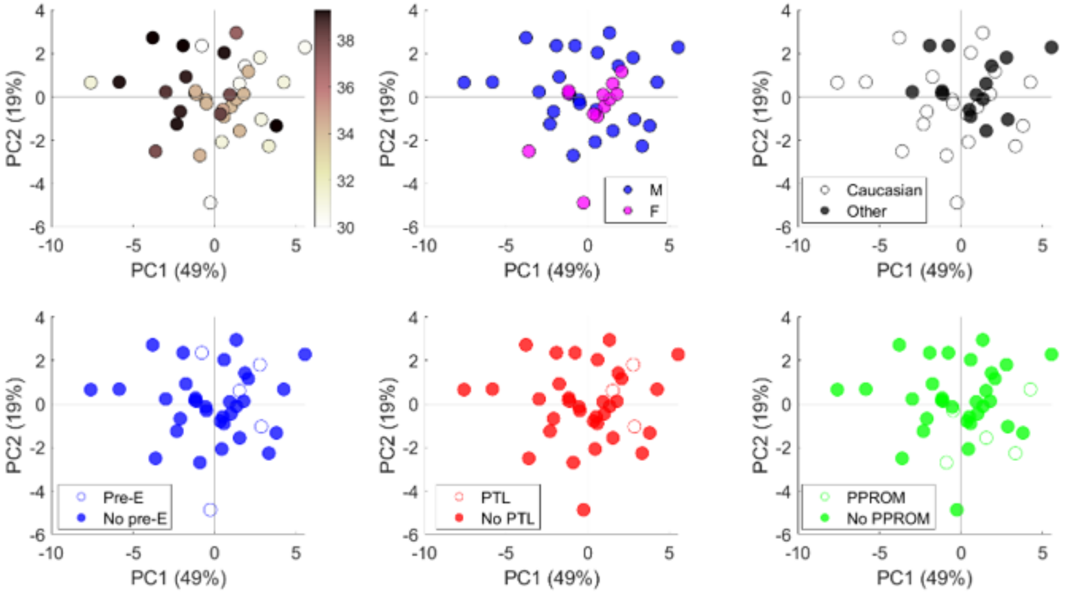


***Supplemental Figure S5. The indication of preterm delivery does not affect placental Fc receptor expression.***

PCA scores plots of patients who delivered prematurely (<37 weeks) in Cohort 2 colored by gestational age, fetal sex, maternal race, or indications of preterm birth (preeclampsia, Pre-E, preterm labor, PTL, or premature preterm rupture of membranes, PRROM).

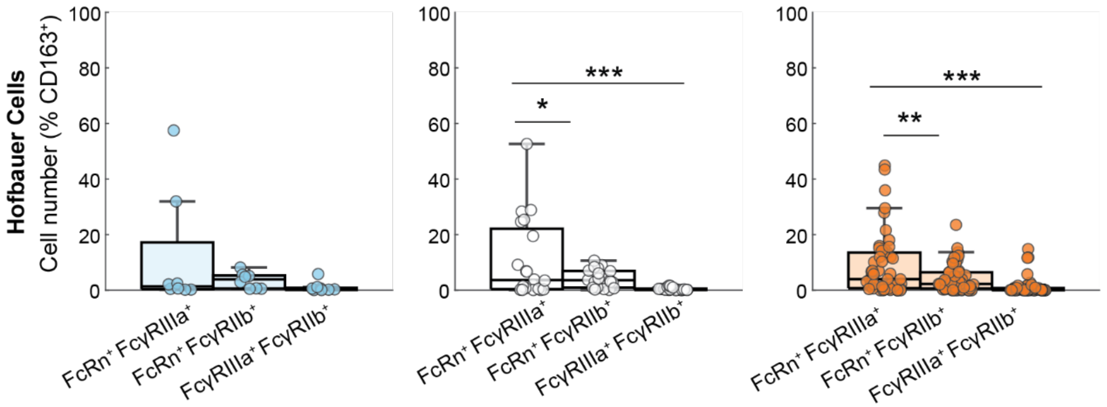


***Supplemental Figure S6. Pairwise colocalization analysis of Fc receptors expressed by Hofbauer cells.***

Boxplots show pairwise colocalizations of Fc receptors on Hofbauer cells, expressed as the average percentage of total Hofbauer cells in each image from each patient. Patients from trimesters are shown on separate axes (blue, 1$^{st}$ trimester; white, 2$^{nd}$ trimester; orange, 3$^{rd}$ trimester). (****p*<0.001, ***p*<0.01, **p*<0.05, ANOVA with Tukey's test for multiple comparisons).

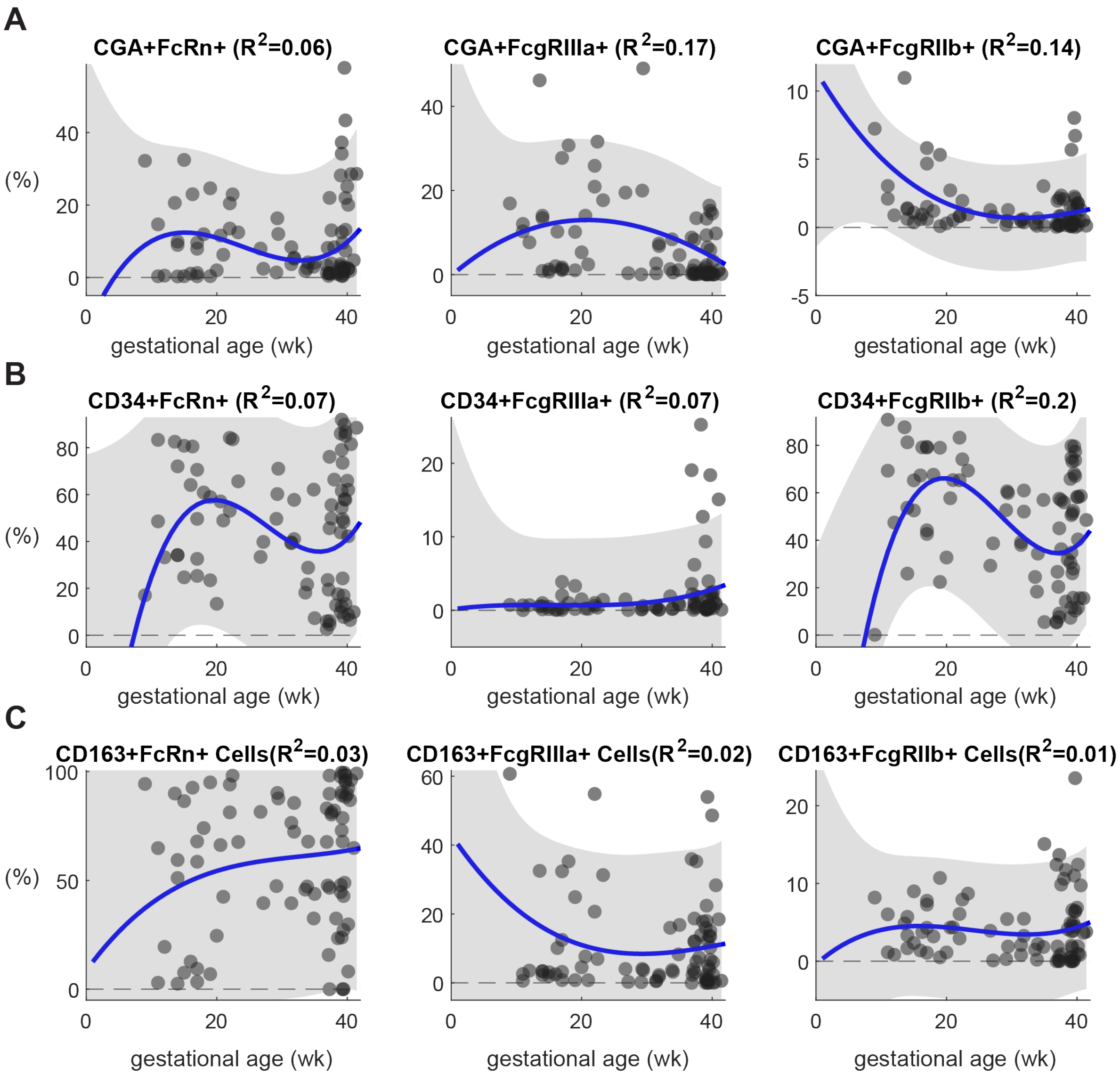


***Supplemental Figure S7. Polynomial regression analysis of cell type-specific Fc receptor expression across gestation in Cohort 1.***

(A-C) Scatter plots showing inferred gestational expression trajectories for each receptor expressed on STBs (A), ECs (B), and HBCs (C) in Cohort 1. Each point represents the average expression frequency from a single patient tissue. The solid lines represent a 3rd-order polynomial curve fit. The gray shaded region denotes 95% confidence intervals. The regression $R^2$ is listed in the top right of each coordinate plane.

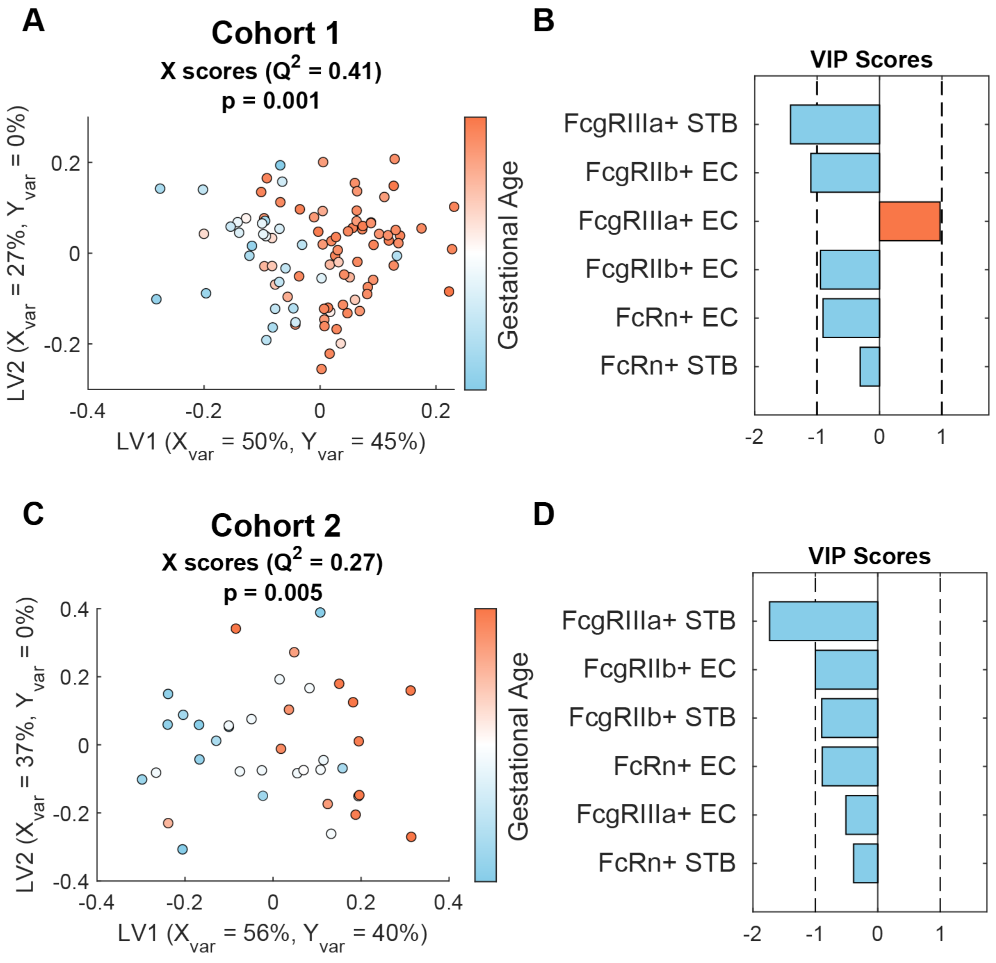


***Supplemental Figure S8. Similar temporal trends in cell type specific Fc receptor expression observed across cohorts.***

OPLSR models trained on Fc receptor expression data from Cohort 1 (A-B) and cohort (C-D). (A,C) X scores plot, where points are colored according to gestational age. Each point represents one patient tissue. $Q^2$ is a metric of the model's prediction performance, and the p-value was determined by permutation testing by comparing $Q^2$ from this model against 1,000 models with randomly shuffled Y-labels. (B,D) VIP scores bar plot. The model trained on data from Cohort 1 (A-B) was used to predict gestational age in Cohort 2, as shown in Figure 5.

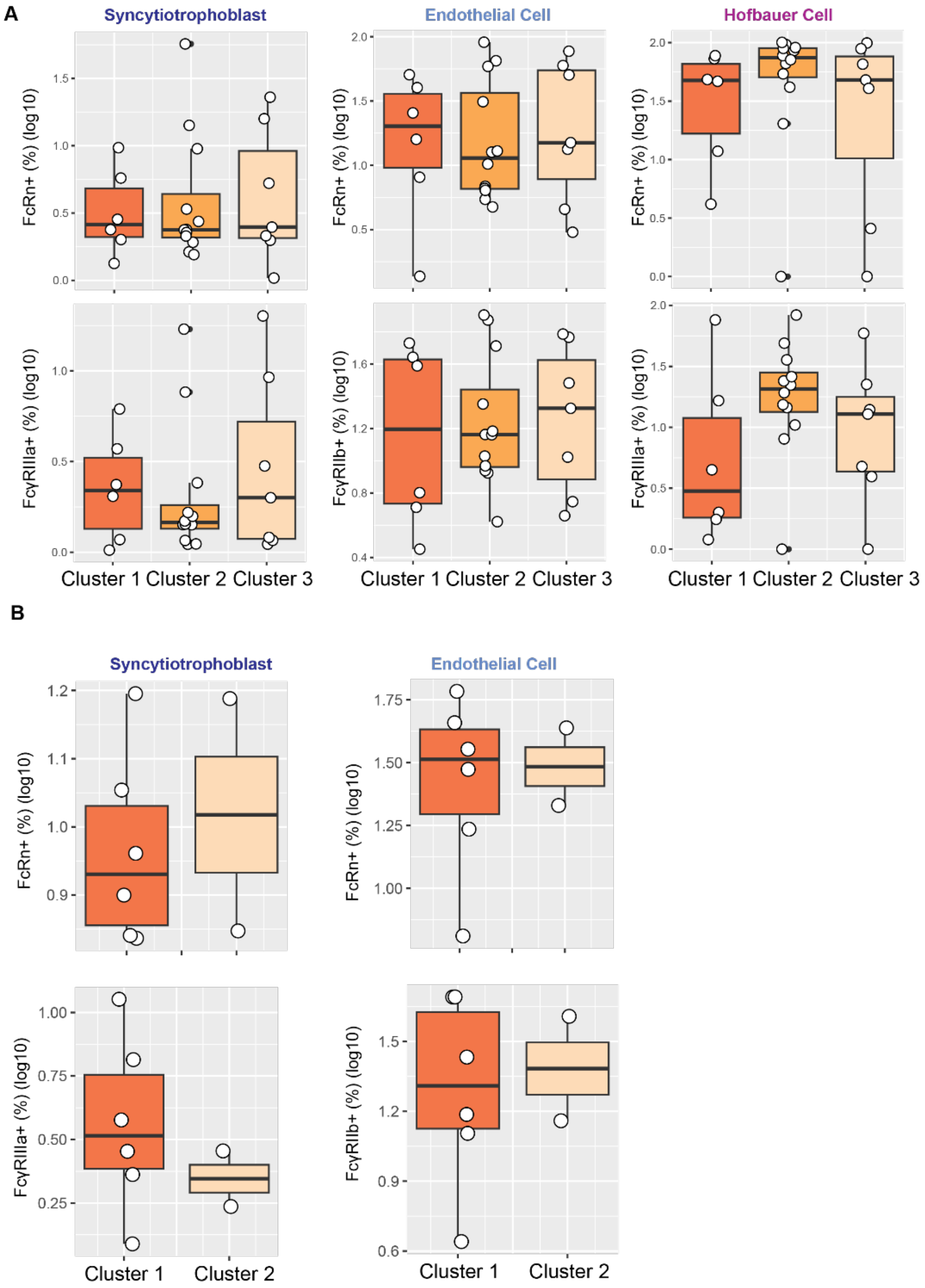


***Supplemental Figure S9. Fc receptor expression across patient clusters.***

(A) Cell type-specific Fc receptor expression levels across clusters in Cohort 1, related to Figure 1. No comparisons were found to be significantly different at ($p<0.1$) (Kruskal-Wallis test with Dunn's test for multiple comparisons). (B) Cell type-specific Fc receptor expression levels across clusters in Cohort 2, related to Figure 1. No comparisons were found to be significantly different at ($p<0.1$) (Kruskal-Wallis test with Dunn's test for multiple comparisons). Only clusters 1 and 3 are shown because no patients in cluster 2 had matched placental tissue and plasma available.

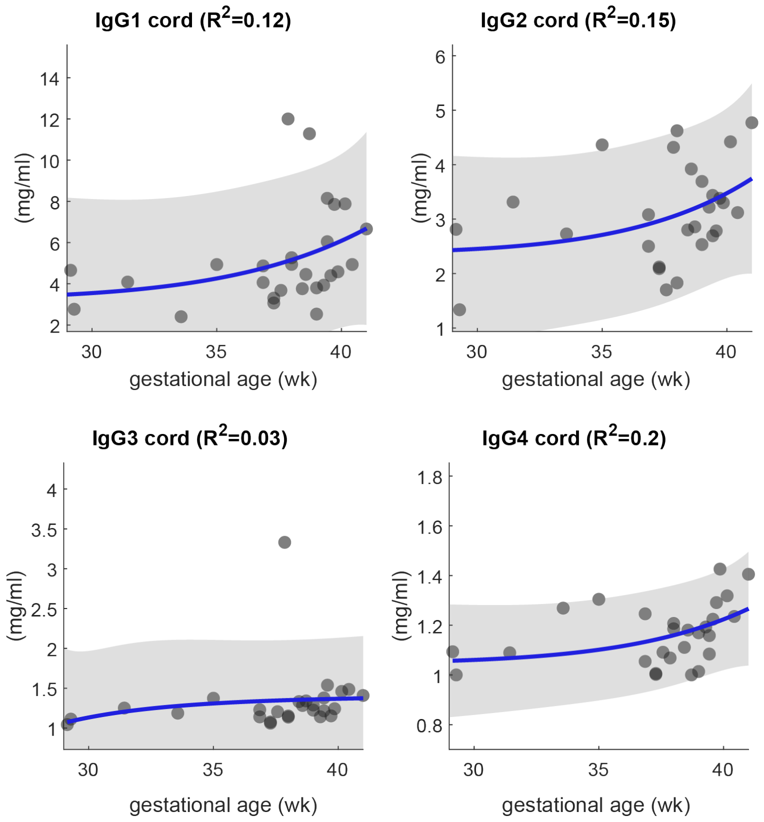


***Supplemental Figure S10. IgG transfer is dynamically regulated across third trimester.***

Scatter plots show the concentration of umbilical cord IgG subclasses as a function of gestational age in Cohort 1. The solid lines are exponential regressions with 95% confidence intervals shaded in gray. Regression $R^2$ are shown in each figure title.

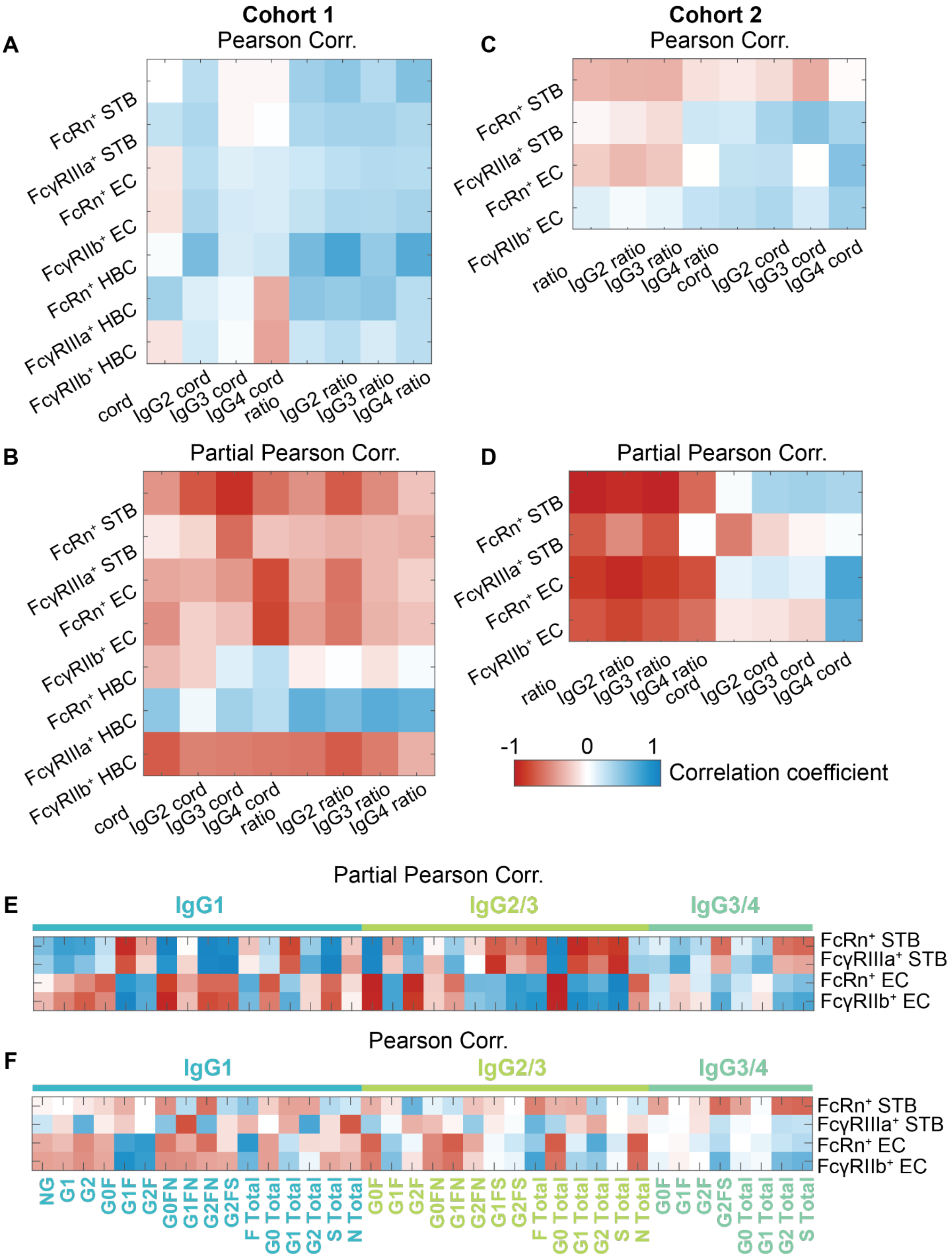
Cohort 1
Pearson Corr.
A
FcRn⁺ STB
FcγRIIIa⁺ STB
FcRn⁺ EC
FcγRIIb⁺ EC
FcRn⁺ HBC
FcγRIIIa⁺ HBC
FcγRIIb⁺ HBC
cord
IgG2 cord
IgG3 cord
IgG4 cord
ratio
IgG2 ratio
IgG3 ratio
IgG4 ratio
Cohort 2
Pearson Corr.
C
B
Partial Pearson Corr.
D
Partial Pearson Corr.
-1
0
1
Correlation coefficient
E
Partial Pearson Corr.
IgG1
IgG2/3
IgG3/4
F
Pearson Corr.
NG
G1
G2
G0F
G1F
G2F
G0FN
G1FN
G2FN
G2FS
F Total
G0 Total
G1 Total
G2 Total
S Total
N Total
G1FS

***Supplemental Figure S11. Controlling for clinical covariates reveals positive associations between Fc receptor expression frequencies and IgG subclass transfer efficiency.***

(A-D) Heatmaps show Pearson correlation coefficients (A,C) and partial Pearson correlation coefficients (B,D) between cell type-specific Fc receptor expression frequencies and IgG subclass cord concentrations or cord:maternal ratios after controlling for clinical covariates used for clustering analysis in Figure 1. Data from Cohort 1 are shown in (A,B) and data from Cohort 2 are shown in (C,D). (E,F) Heatmaps show Pearson correlation coefficients (E) and partial Pearson correlation coefficients (F) between cell type-specific Fc receptor expression frequencies and subclass-specific glycoforms after controlling for clinical covariates used for clustering analysis in Figure 1. The colored bar denotes the subclass-specific glycoforms. Glycoforms were omitted from the heatmap if they fell below the limit of detection, resulting in NA correlation values with all Fc receptors.

***Supplemental Table 1. Intact glycopeptide/non-glycosylated peptide masses of IgG subclasses Fc conservative N-glycosylation.***

| Glycoform* | IgG1 (EEQYN*STYR) $[M+H]^+$ | IgG2/3 (EEQFN*STFR) $[M+H]^+$ | IgG3/4 (EEQFN*STYR/EEQYN*STFR) $[M+H]^+$ |
|---|---|---|---|
| NG | 1189.5120 | 1157.5222 | 1173.5171 |
| G0 | 2487.9879 | 2455.9981 | 2471.9930 |
| G1 | 2650.0407 | 2618.0509 | 2634.0458 |
| G2 | 2812.0936 | 2780.1038 | 2796.0987 |
| G0N | 2691.0673 | 2659.0775 | 2675.0724 |
| G1N | 2853.1201 | 2821.1303 | 2837.1252 |
| G2N | 3015.1729 | 2983.1831 | 2999.1780 |
| G1S | 2941.1362 | 2909.1464 | 2925.1413 |
| G2S | 3103.1890 | 3071.1992 | 3087.1941 |
| G1NS | 3144.2155 | 3112.2257 | 3128.2206 |
| G2NS | 3306.2684 | 3274.2786 | 3290.2734 |
| G0F | 2634.0458 | 2602.0560 | 2618.0509 |
| G1F | 2796.0987 | 2764.1089 | 2780.1038 |
| G2F | 2958.1515 | 2926.1617 | 2942.1566 |
| G0FN | 2837.1252 | 2805.1354 | 2821.1303 |
| G1FN | 2999.1780 | 2967.1882 | 2983.1831 |
| G2FN | 3161.2309 | 3129.2411 | 3145.2360 |
| G1FS | 3087.1941 | 3055.2043 | 3071.1992 |
| G2FS | 3249.2469 | 3217.2571 | 3233.2520 |
| G1FNS | 3290.2734 | 3258.2836 | 3274.2785 |

* There is no consortium enforcing a standardized nomenclature for IgG conservative Fc N-glycosylation[7]. The composition shorthand codes in this table are NG= non-glycosylated, G0=agalactosylated, G1/2=mono- and double- galactosylated, F=core fucosylated, N=bisecting *N*-acetylglucosaminated, S=monosialylated[4].

***Supplemental Table 2. Retention time ranges for glycopeptides/non-glycosylated peptides from all IgG subclasses.***

| Subclass | Neutral RT (min) | Acidic RT (min) | NG RT (min) |
|---|---|---|---|
| IgG1 (EEQYN*STYR) | 9.6-10.8 | 14.1-15.7 | 10.3 |
| IgG2/3a (EEQFN*STFR) | 17.9-19.8 | 23.5-25.2 | 19.8 |
| IgG3b/4 (EEQFN*STYR/EEQYN*STFR) | 13.2-14.1 | 18.5-18.9 | Not Detected |
